# Molecular mechanism of redox regulation of the alpha-carboxysomal carbonic anhydrase CsoSCA

**DOI:** 10.64898/2026.04.02.716132

**Authors:** Nikoleta Vogiatzi, Guillaume Gaullier, Joel Leufstadius, Tor Andersson, Thomas Scherbauer, Cecilia Blikstad

## Abstract

Carboxysomes are protein-based organelles that form the core of the bacterial CO_2_-concentrating mechanism (CCM) by elevating CO_2_ levels around Rubisco. They encapsulate Rubisco and carbonic anhydrase (CA) within a protein shell that, after closure, excludes cytosolic reductants. Because cytosolic CA activity would short-circuit the CCM, CA activity must be confined to the carboxysome, yet how α-carboxysomes achieve this has remained unknown. Here we show that CsoSCA, the α-carboxysomal CA, is redox-regulated: inactive under reducing conditions, active under oxidizing. This regulation is mediated by a conserved vicinal cysteine pair distal from the active site. CryoEM structures of *Halothiobacillus neapolitanus* CsoSCA under active and inactive conditions, and of an inactive cysteine variant, reveal that redox conditions modulate global conformational dynamics that reorganize the active site for catalysis. These findings advance the understanding of α-carboxysome regulation and couple CsoSCA activation to lumenal oxidation during carboxysome maturation.

## Introduction

Carboxysomes are protein-based bacterial organelles that encapsulate the key enzymes for CO_2_ fixation - Ribulose-1,5-bisphosphate carboxylase/oxygenase (Rubisco) and carbonic anhydrase (CA) - within a protein shell (1, 2). Together with energy-coupled inorganic carbon transporters, carboxysomes form the bacterial CO_2_ concentrating mechanism (CCM) (3, 4). This mechanism, present in all cyanobacteria and many proteobacteria, functions to overcome Rubisco’s modest turnover number and its inability to distinguish between CO_2_ and the competing off-target substrate O_2_ (5, 6). The inorganic transporters actively accumulate HCO_3_^-^ in the cytosol, which then diffuses through the carboxysome shell. Inside, the carboxysomal CA rapidly equilibrates HCO_3_^-^ to CO_2_ producing a locally high CO_2_ concentration around Rubisco. This saturates Rubisco’s active sites with CO_2_, competitively inhibiting oxygenation, and thereby ensures overall carbon assimilation efficiency (7, 8).

Two types of carboxysomes have emerged through convergent evolution: the α-type, predominantly found in oceanic cyanobacteria and proteobacteria, and the β-type, typically present in freshwater cyanobacteria (1, 2, 9, 10). Both types share the same icosahedral structure and are built up by hexameric and pentameric shell proteins, one or two scaffolding proteins, and the encapsulated enzymes Rubisco and CA. Carboxysomes self-assemble through a condensate-driven mechanism that coordinates the formation of both cargo and shell (11, 12). Once fully assembled, the shell is thought to act as a barrier which excludes cellular reductants, generating an oxidizing interior distinct from the reducing cytosol (13–15).

CAs catalyze the rapid interconversion between CO_2_ and HCO_3_^-^ In photosynthesis, their primary role is often to supply Rubisco with CO_2_ (16). CAs are a diverse class of enzymes. To date, there are eight characterized families (α, β, γ, δ, ζ, η, θ and ι) which have evolved convergently and share little to no sequence or structural similarity (17, 18). Some CAs exhibit a catalytic efficiency (*k*_cat_/*K*_M_) reaching 10^9^ M^-1^s^-1^, making them among the fastest known enzymes. However, the carboxysomal CAs characterized so far display more modest activity, with *k*_cat_/*K*_M_ values in the range of 10^6^-10^7^ M^-1^s^-1^ (19–21). β-carboxysomes encapsulate either a β-CA, named CcaA (20, 22), or an active γ-CA domain on the scaffolding protein CcmM (19, 23). The vast majority of α-carboxysomes encapsulate the β-CA CsoSCA (21, 24–27), although, a recent study identified a novel ι-CA in α-carboxysomes from a sulfur-oxidizing alkaliphilic γ-proteobacterium (28).

CsoSCA consists of three distinct domains (Fig. 1A) (26). The N-terminal domain includes an intrinsically disordered peptide essential for carboxysome encapsulation (24), followed by a folded domain unique to CsoSCA and recently shown to mediate oligomerization (29, 30). The middle catalytic domain contains the Zn^2+^-binding site and essential catalytic residues (26), including the catalytic dyad typical of CAs. The C-terminal domain likely arose from an ancient gene duplication of the catalytic domain but lacks the active site residues. In the cyanobacterial lineage, this domain harbors an RuBP-binding site involved in allosteric activation (29), however, in proteobacteria its function remains unknown.

**Figure 1:**
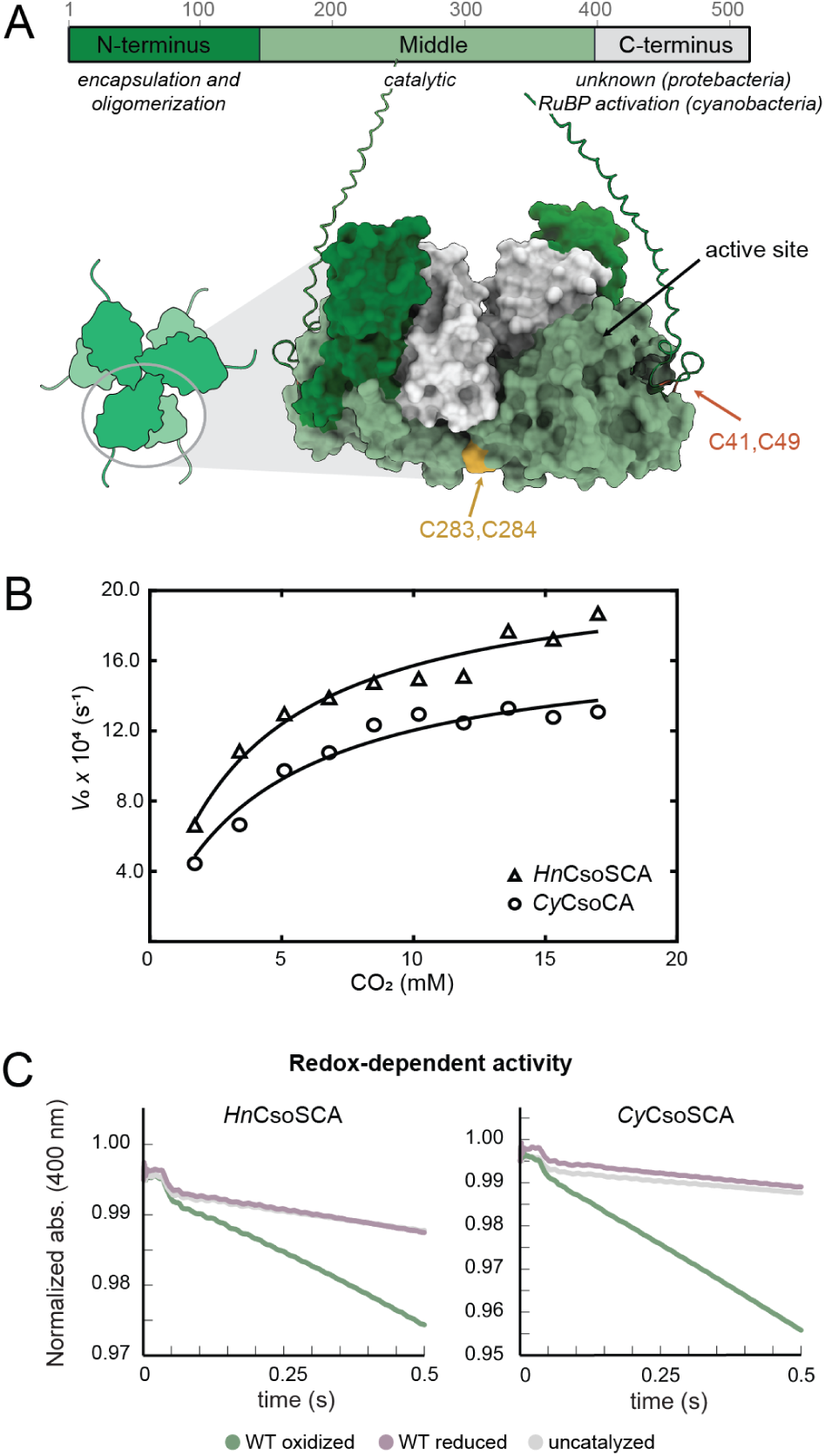
CsoSCA is redox-regulated across species. (A) Schematic of *Hn*CsoSCA’s domain structure and its oligomeric organization as a trimer of dimers, alongside an AlphaFold2 model of a dimer illustrating the three domains. The N-terminal domain mediates encapsulation and oligomerization and consists of an intrinsically disordered N-terminus followed by a folded domain. The middle domain contains the active site. The C-terminal domain has an unknown function in *Hn*CsoSCA but is involved in allosteric activation in *Cy*CsoSCA. Two cysteine pairs were identified to be within compatible distance for disulfide bond formation: Cys283,Cys284 (yellow) and Cys41,Cys49 (orange). Residue numbering corresponds to *Hn*CsoSCA. (B) Saturation curves of *Hn*CsoSCA (triangles) and *Cy*CsoSCA (circles) catalyzed CO_2_ hydration at pH 8. See table 1 for kinetic constants. (C) Activity measurements of *Hn*CsoSCA and *Cy*CsoSCA in oxidizing and reducing conditions, showing that both enzymes are only active in oxidizing conditions and inactive when reduced.

**Table 1:** Steady-state kinetic parameters for CsoSCA-catalyzed CO_2_ hydration. Measurements were performed at 25 °C using the HEPES-phenol red buffer-indicator pair at pH 8.0, at a wavelength of 558 nm.

| Protein | Condition | $k_{\text{cat}}$ (s<sup>-1</sup>) | $K_M$ (mM) | $k_{\text{cat}}/K_M$ (M<sup>-1</sup>s<sup>-1</sup>) |
| --- | --- | --- | --- | --- |
| *HnCsoSCA* | Oxidizing | $2.1 \times 10^4 \pm 9.6 \times 10^2$ | $3.7 \pm 0.6$ | $5.8 \times 10^6 \pm 6.5 \times 10^2$ |
| CyCsoSCA | Oxidizing | $1.7 \times 10^4 \pm 9.9 \times 10^2$ | $4.3 \pm 0.7$ | $4.0 \times 10^6 \pm 5.0 \times 10^2$ |

Knock-out experiments have shown that CA activity in the carboxysome is an absolute requirement for the CCM to function (24, 31, 32). Alongside this, early work by Price and Badger demonstrated that heterologous expression of CA in the cyanobacterial cytosol leads to a high CO_2_-requiring phenotype (33), supporting the hypothesis that cytosolic CA activity short-circuits the CCM by equilibrating accumulated HCO_3_^-^ to CO_2_ an uncharged molecule which can freely diffuse out of the cell (2). Thus, efficient encapsulation and tight regulation of carboxysomal CAs are critical for maintaining cellular function. We recently detailed how CsoSCA is efficiently encapsulated into α-carboxysomes via direct interactions with Rubisco (24). While the regulatory mechanism for the γ-class CA CcmM in β-carboxysomes is understood (23), how CsoSCA remains inactive in the cytosol and activates upon encapsulation into functional α-carboxysomes remains unknown.

Here, we measured steady-state enzyme kinetics on CsoSCA using the stopped-flow based Khalifha/pH indicator assay and established that CsoSCA is active under oxidizing conditions and inactive under reducing conditions. We pin-point a conserved vicinal disulfide distal from the active site as the regulatory switch and show that this regulation is conserved across species, occurring in orthologs from both proteobacteria and cyanobacteria. We further determined single-particle cryo-electron microscopy (cryo-EM) structures of CsoSCA from the model organism *Halothiobacillus neapolitanus c2* in oxidizing (2.15 Å resolution) and reducing (2.12 Å resolution) conditions, as well as of an inactive cysteine variant (2.18 Å resolution). The structures show that oxidizing conditions enable CsoSCA to adopt a globally closed conformation that correctly positions key active site residues for catalysis. In contrast, reduction favors a globally open conformation that disrupts essential hydrogen bonding networks and thereby abolishes activity. Combined, our findings reveal the role of protein dynamics in the function of CsoSCA and define the molecular mechanism governing α-carboxysomal CA regulation.

## Results

### Kinetic characterisation of *Hn*CsoSCA and *Cy*CsoSCA

To characterize the kinetic behavior and identify the regulatory mechanism of CsoSCA in α-carboxysomes, we studied two homologs of the enzyme: *Hn*CsoSCA from *H. neapolitanus*, a γ-proteobacterial model organism, and *Cy*CsoSCA from *Cyanobium* sp. PCC 7001, a cyanobacterial ortholog. Initial purification attempts showed that full-length CsoSCA from both species readily aggregate, preventing us from obtaining quantities sufficient for kinetic measurements. Fusing an MBP solubility tag to the N-terminus yielded high quantities of stable and pure MBP-*Hn*CsoSCA and MBP-*Cy*CsoSCA. Attempts to cleave off the MBP-tag resulted in severe precipitation. Further, we compared His- and Strep-tagged constructs: the His-tagged proteins showed 10-fold lower activity as compared to Strep-tagged, suggesting that the His-tag interferes with the active site and impairs the activity. Based on these findings, we proceeded with N-terminal Strep-MBP fusion proteins to ensure high concentration of well behaved CsoSCA needed for activity assays.

To measure steady-state kinetics of CsoSCA-catalyzed CO_2_ hydration, we used the stopped-flow based Khalifah/pH indicator assay (34, 35). Both MBP-*Hn*CsoSCA and MBP-*Cy*CsoSCA were found to be catalytically active, and kinetic constants were determined by measuring saturation curves in the range of 1.7 - 17 mM CO_2_. In both instances, the turnover number (*k*_cat_) was determined to ∼2 x 10^4^ s^-1^ and the Michaelis-Menten constant (*K*_M_) to ∼4 mM (Fig. 1B, Table 1). This corresponds to catalytic efficiencies (*k*_cat_/*K*_M_) in the order of 10^6^ M^-1^s^-1^, which is comparable to published values for untagged *Hn*CsoSCA (21), concluding that the N-terminal Strep-MBP tag does not affect activity.

### The enzymatic activity of CsoSCA is redox-regulated

From kinetics data and crystal structures, the γ-class CA in β-carboxysomes, CcmM, from *Thermosynechococcus vestitus BP-1* (formerly *T. elongatus BP-1*) was shown to be redox-regulated (23). Under oxidizing conditions, a disulfide is formed in a loop distant from the active site, activating CcmM, while reducing conditions disrupt this disulfide and render the enzyme inactive. Similarly, reducing agents have been suggested to inhibit *Hn*CsoSCA activity (21, 27), hinting at a similar redox-dependent mechanism of CA regulation in α-carboxysomes despite the fact that the two enzymes, CcmM and CsoSCA, are structurally distinct and belong to different CA-classes. To validate the hypothesis that CsoSCA’s activity is regulated by its redox environment, we measured specific activity at 17 mM CO_2_ under oxidizing and reducing conditions for both *Hn*CsoSCA and *Cy*CsoSCA. Both orthologs were active in oxidizing conditions but no enzymatic activity could be detected when the enzymes were reduced prior to the measurements (Fig. 1C, SI Fig. 1A). Repeating the measurements with a 10-fold higher enzyme concentration verified the absence of physiologically relevant activity under reducing conditions (SI Fig. 2A).

### A conserved vicinal cysteine pair acts as a regulatory switch

Structural and sequence analysis identified two pairs of cysteines positioned to allow formation of disulfide bonds: (I) a potential disulfide formed by two vicinal cysteines located ∼35 Å from the active site, present in both *Hn*CsoSCA (Cys283,Cys284) and *Cy*CsoSCA (Cys353,Cys354), and (II) a potential disulfide in the predicted disordered N-terminus of *Hn*CsoSCA (Cys41,Cys49) (Fig. 1A). No additional candidates were found in *Cy*CsoSCA. To assess their role in the observed redox regulation, these cysteine pairs were mutated to alanines and enzyme variants tested for activity. The *Hn*CsoSCA_C283A,C284A_, and *Cy*CsoSCA_C353A,C354A_ variants showed no detectable activity, whereas *Hn*CsoSCA_C41A,C49A_ retained WT-like activity (Fig. 2A and B, SI Fig. 1B and C). To ensure that *Hn*CsoSCA_C283A,C284A_ and *Cy*CsoSCA_C353A,C354A_ were truly inactive, their activity was tested with enzyme concentrations 10-fold higher than for WT and no activity could still be detected (SI Fig. 2B), establishing the role of this cysteine pair as the redox switch.

**Figure 2:**
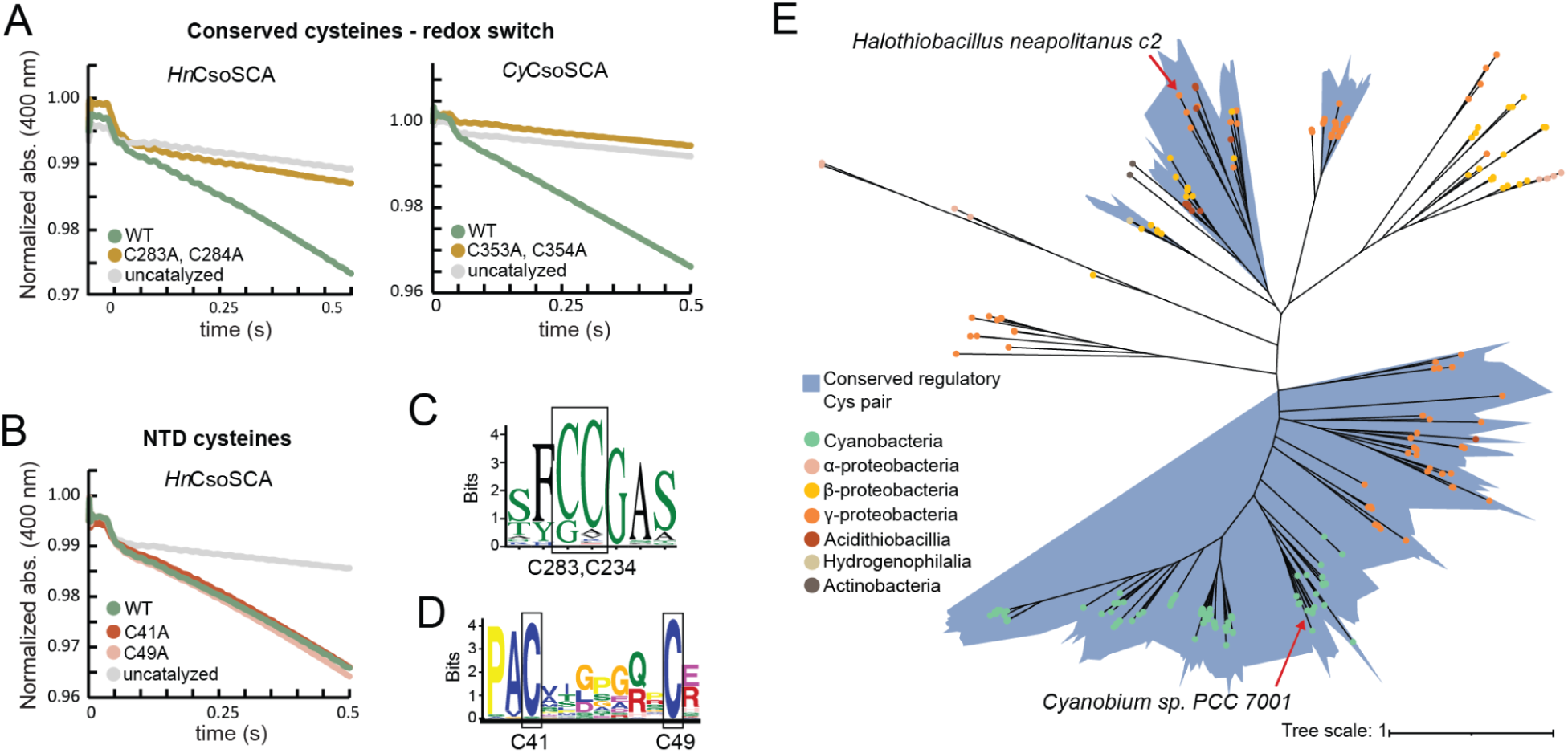
A conserved vicinal cysteine pair acts as a redox switch. (A) Activity measurements of *Hn*CsoSCA_C283A,C284A_ and *Cy*CsoSCA_C353A,C354A_ showing that these alanine mutants are inactive and suggesting that the conserved cysteine pair acts as a regulatory redox switch. (B) Activity measurements of *Hn*CsoSCA_C41A,C49A_ showing that the activity is not affected by disrupting the disulfide in the N-terminal disordered region of *Hn*CsoCSA. (C) Multiple sequence alignment of 222 CsoSCA sequences visualized as a sequence logo showing strong conservation of the regulatory cysteine pair. (D) Motif discovery of the intrinsically disordered N-terminus using MEME analysis. 42 out of 222 sequences contain the PACxxxxxxxC motif. Residue numbering corresponds to *Hn*CsoSCA. (E) Unrooted maximum-likelihood phylogenetic tree of CsoSCA. Clades containing the conserved regulatory cysteine pair are highlighted in blue. A version of this tree with all species annotated is shown in SI Fig. 3.

Re-analysis of 222 CsoSCA sequences from Blikstad *et al*. (24) shows that the identified regulatory cysteine pair is conserved in ∼80% of orthologs, suggesting a conserved regulatory mechanism (Fig. 2C). This vicinal cysteine pair occurs in all cyanobacterial sequences and in ∼65% of the proteobacterial sequences (Fig. 2E, SI Fig. 3). From sequence alignment and structural analysis, we could not identify any other cysteines that could clearly have this function in the three phylogenetic clades that lack the regulatory cysteine pair. In summary, these results suggest that CsoSCA is inactive in the reducing cytosolic environment and activates once encapsulated in the oxidizing milieu of fully closed and matured carboxysomes. Phylogenetic analysis and targeted mutagenesis indicate that this regulation mechanism is conserved across phyla and is mediated by a conserved vicinal cysteine pair located strikingly far away from the active site.

### *Hn*CsoSCA is organized as a hexamer regardless of redox conditions

To further elucidate the molecular mechanism of CsoSCA’s redox regulation, we determined the structure of *Hn*CsoSCA under both oxidizing and reducing conditions. Using the same protein preparations as in the activity assays, MBP-*Hn*CsoSCA was vitrified in oxidizing buffer and in reducing buffer, and cryo-EM data were collected on the enzyme in these two conditions. Additionally, cryo-EM data were collected on the inactive *Hn*CsoSCA_C283A,C284A_ variant in oxidizing buffer.

In all three reconstructions, neither the MBP-tag nor the disordered N-terminus are resolved, and *Hn*CsoSCA forms a homo-hexamer organized as a trimer of dimers and was therefore refined with D3 symmetry imposed (oxidizing-hexamer: PDB 9SKR, 2.13 Å resolution; reducing-hexamer: PDB 9SKU, 2.06 Å resolution; *Hn*CsoSCA_C283A,C284A_ hexamer: PDB 9SKX, 2.08 Å resolution) (Fig. 3A, SI Fig. 4-6). This quaternary structure is in contrast to a previous crystal structure of a Y92H variant which had suggested *Hn*CsoSCA to be dimeric (PDB 2FGY (26). However, a hexameric assembly is supported in our previous work by size exclusion chromatography (24) and by recently published WT structures of both *Hn*CsoSCA (PDB 9G4T (30) and *Cy*CsoSCA (PDB 8THM (29). Importantly, the fact that the hexameric quaternary structure is unaffected by redox conditions or mutation of the redox-regulating cysteines demonstrates that trimerization does not play a regulatory role (Fig. 3B).

**Figure 3:**
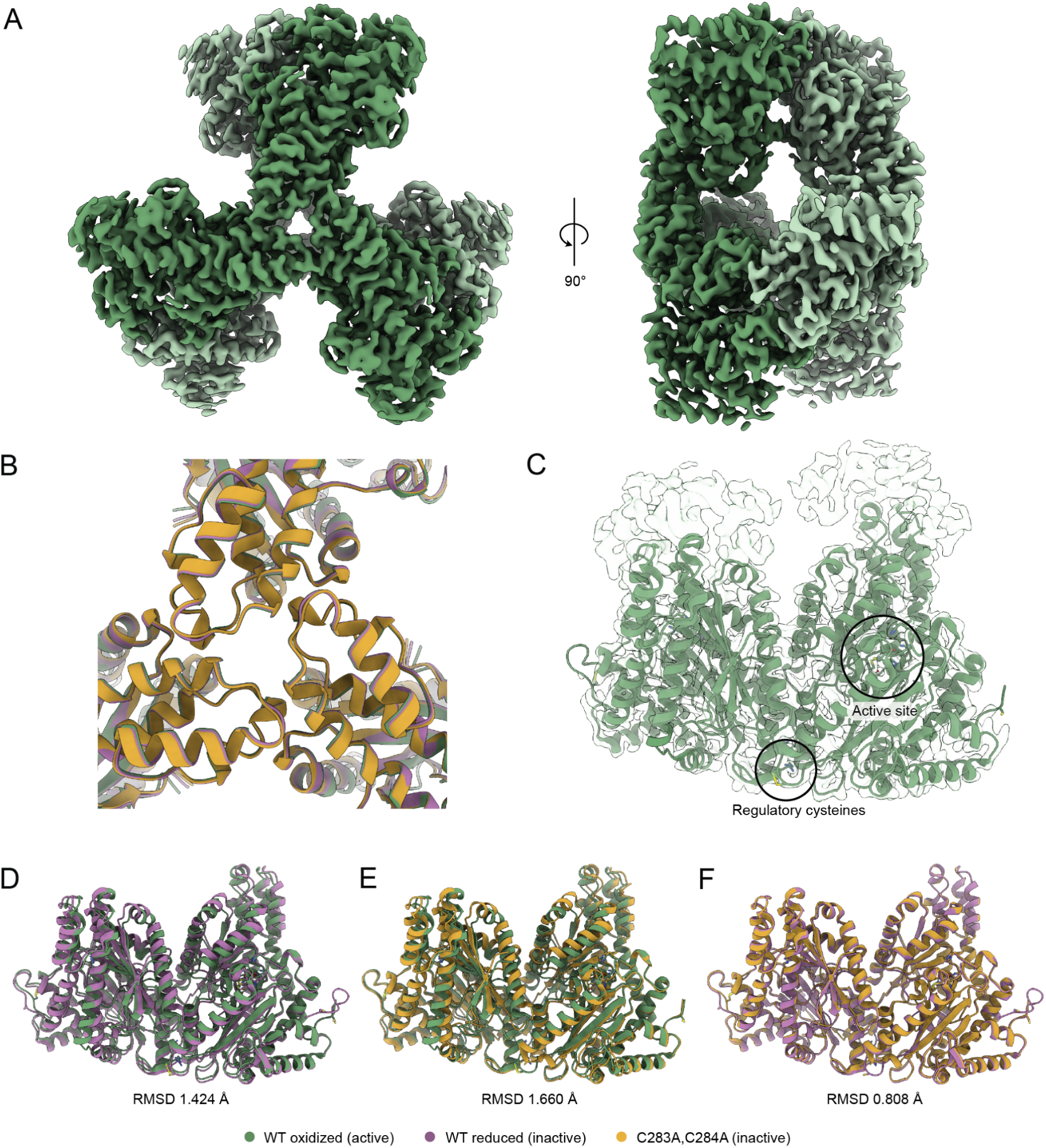
Structures of WT HnCsoCA in oxidizing and reducing conditions and of the inactive C283A,C284A variant. (A) Cryo-EM map of *Hn*CsoSCA in oxidizing conditions showing that it is organized as a homohexamer: a trimer of dimers. The two protomers in a dimer are colored in different shades. (B) Close-up of the trimerization interface. Superimposition of WT oxidized, WT reduced and C283A,C284A variant showing that the oligomerization is unaffected by redox condition and mutation. (C) Atomic model of the *Hn*CsoSCA oxidized-closed dimer with the corresponding cryo-EM map overlaid as a translucent surface. The locations of the active site and regulatory cysteines are marked with circles. (D-F) Pairwise superimpositions of the dimer atomic models showing differences in global conformation between: (D) oxidized-closed vs reduced-open, (E) oxidized-closed vs C283A,C284A-open and (E) reduced-open vs C283A,C284A. RMDS values from these superimpositions are indicated. WT oxidized structures are shown in green, WT reduced in purple and the C283A,C284A variant in yellow.

**Figure 4:**
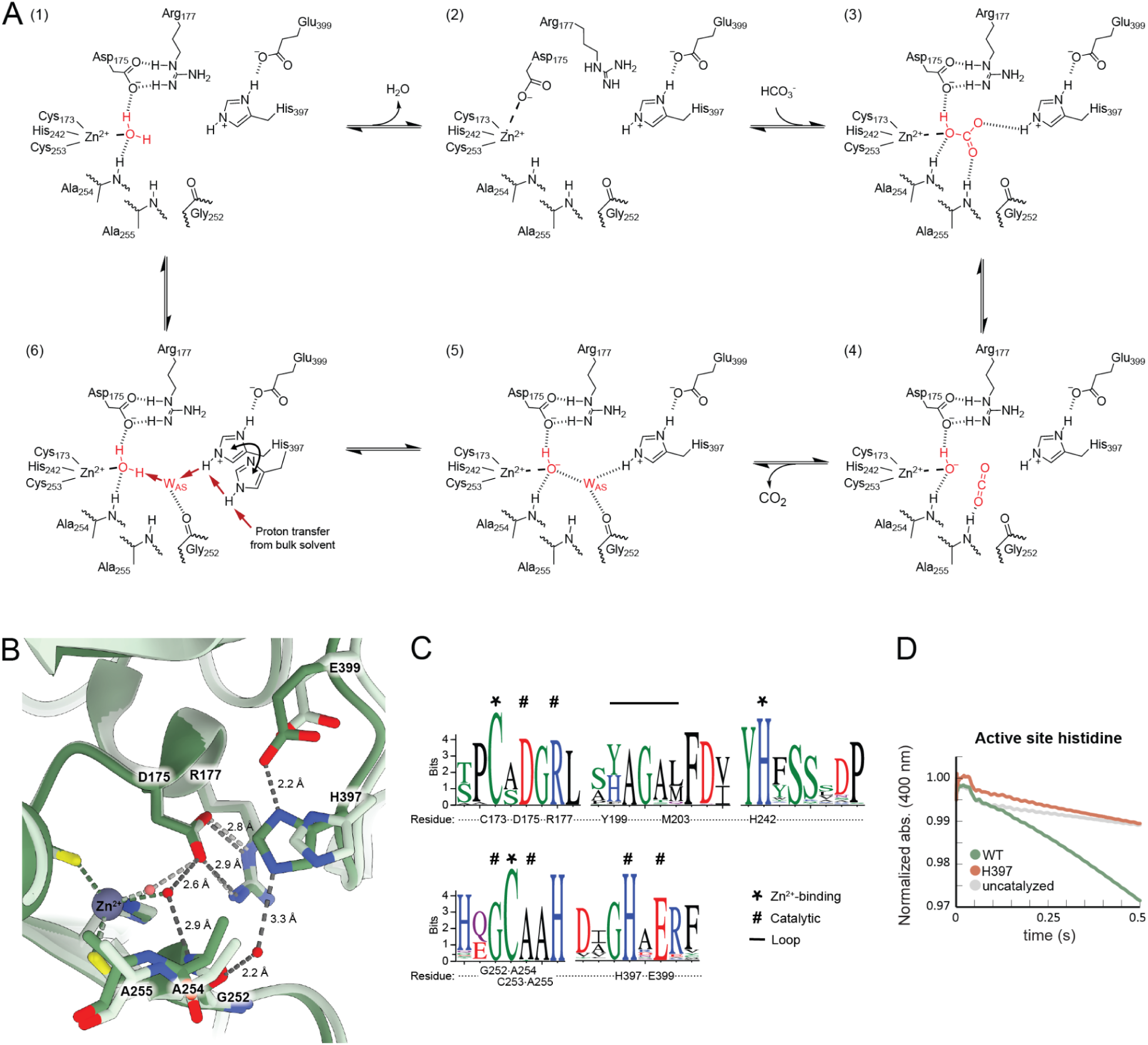
Proposed catalytic mechanism of CsoSCA catalyzed HCO_3_^-^ dehydration. (A) Proposed reaction scheme. The Zn^2+^-coordinated water (1) gets displaced by Asp175 (2). HCO_3_^-^ enters the active site and coordinates between the Zn^2+^, Ala254, Ala255 and His397 (3). HCO_3_^-^ decomposes to CO_2_ and OH^-^ (4) and CO_2_ diffuses out of the active site (5). The Zn^2+^-bound OH^-^ gets protonated by the active site water (W_AS_) (6) restoring the active site. We propose that this proton transfer is mediated by His397 that carries a proton from the bulk solvent to W_AS_ when it shifts from its far to its near position. We cannot distinguish in which steps W_AS_ is present and it is therefore only depicted in steps in which it has a functional role. (B) Close-up view of the active site comparing the distances and hydrogen bond network involved in catalysis between the oxidized-closed (green) and oxidized-open structures (light green). Hydrogen bonds are colored in dark grey for oxidized-closed and light grey for oxidized-open. Water molecules are colored in red for oxidized-closed and light red for oxidized-open. Indicated distances are those observed in the oxidized-closed structure. (C) Multiple sequence alignment visualized as sequence logo showing the conservation of active site residues involved in Zn^2+^-coordination, catalysis and forming the 199-203 loop. (D) Activity measurements of *Hn*CsoSCA_H397A_ demonstrating loss of activity confirming the essential catalytic role for His397.

### *Hn*CsoSCA exists in an equilibrium between different global conformations

The regulatory cysteine pair is located far from the active site, ∼35 Å from the catalytic Zn²⁺ in the same protomer, in a loop at the dimer interface. We therefore suspected that the regulation involves long-range effects like conformational changes and/or differences in protein dynamics. To reveal such effects, we performed 3D variability analysis (3DVA) without symmetry on each set of particles. This revealed that the symmetry-averaged hexameric reconstructions of WT *Hn*CsoSCA, under both oxidizing and reducing conditions, represent a mixture of conformational states, and that the main component of variability is a flexing motion around the dimer axis (Movie 1 and 2). The inactive C283A,C284A variant, on the other hand, shows some dynamics but with smaller amplitude and with this flexing around the dimer axis no longer contributing to the first component of variability (Movie 3), suggesting that this variant is more conformationally homogeneous. To sort these different conformations, we applied symmetry expansion to define the dimer as the unit of interest and subjected all dimers in each dataset to 3D classification without alignment followed by local refinement. Atomic models were then built and refined against each of these dimer reconstructions (Fig. 3C, SI Fig. 7).

For the WT enzyme under both redox conditions, 3D classification indeed identified two populations of dimers with distinct global conformations: one more globally “closed” and the other more globally “open” around the dimer axis. From here on, we refer to these different global conformations as “closed” and “open” (Table 2). Applying the same classification to the C283A,C284A variant yielded two structures with only subtle local differences (RMSD 0.219 Å, SI Fig. 6 and 8C), both having the global open conformation. For *Hn*CsoSCA_C283A,C284A_, we will therefore only describe the structure assigned PDB code 9SKY which has better defined density around the active site. To identify each dimer structure without ambiguity, from here on we refer to them by the combination of redox condition or mutation, and global conformation: oxidized-closed (PDB 9SKS), oxidized-open (PDB 9SKT), reduced-closed (PDB 9SKW), reduced-open (PDB 9SKV) and C283A,C284A-open (PDB 9SKY, 9SKZ) (Table 2).

**Table 2:** Overview of *Hn*CsoSCA dimer structures. Global and local features that differ markedly between structures are listed. A and B refer to chain identifiers and numbers refer to residue numbers in the deposited PDB entries. For comparison of position of active site residues, distances are measured from the hydrogen bonding residue to the catalytic Zn^2+^. For the active site water, distance is measured to His397.

|  | WT oxidized |  | WT reduced |  | C283A,C284A |
| --- | --- | --- | --- | --- | --- |
| **Activity detected** | Yes |  | No |  | No |
| **PDB code** | 9SKS | 9SKT | 9SKV | 9SKW | 9SKY |
| **Resolution (Å)** | 2.15 | 2.45 | 2.12 | 2.27 | 2.18 |
| **Global conformation** | Closed | Open | Open | Closed | Open |
| **Proportion of dimers in the dataset (%)** | 54.9<sup>a</sup> | 41.0<sup>a</sup> | 87.7 | 12.3 | 45.2<sup>b</sup> |
| **Loop 199-203** | A: Closed<br>B: Closed | A: Open<br>B: Closed | A: Open<br>B: Open | A: Open<br>B: Open | A: Closed<br>B: Closed |
| **His397**<br>**(Distance: Nε2 of His397 to Zn<sup>2+</sup>)** | A: Near (6.7 Å)<br>B: Near (6.8 Å) | A: Far (8.4 Å)<br>B: Far (8.4 Å) | A: Far (8.7 Å)<br>B: Far (8.7 Å) | A: Near (7.0 Å)<br>B: Near (7.5 Å) | A: Far (8.7 Å)<br>B: Far (8.9 Å) |
| **Glu399**<br>**(Distance: Oε2 of Glu399 to Zn<sup>2+</sup>)** | A: Near (8.4 Å)<br>B: Near (8.4 Å) | A: Far (9.7 Å)<br>B: Far (9.8 Å) | A: Far (9.8 Å)<br>B: Far (9.8 Å) | A: Near (8.4 Å)<br>B: Near (8.8 Å) | A: Far (9.9 Å)<br>B: Far (9.9 Å) |
| **Active site H<sub>2</sub>O**<br>**(Nε2 of His397 to H<sub>2</sub>O)** | A:2101 (3.3 Å)<br>B:2110 (3.4 Å) | A: No<br>B: No | A: No<br>B: No | A: No<br>B: No | A: 2149 (4.2 Å)<sup>c</sup><br>B: No |

<sup>a</sup> The remaining 4.1% of dimers are too affected by air-water interface damage to assign their global conformation.
<sup>b</sup> The remaining 5.7% of dimers are too affected by air-water interface damage to assign their class.
<sup>c</sup> This water is not hydrogen bonding to His397 and is located further away from the active site than the water observed in 9SKS.

Superimposing the different structures (Fig. 3D-F, SI Fig. 8) shows that: (I) the globally closed and open conformations are clearly different from each other (e.g. RMSDs: 1.032 Å oxidized-closed vs oxidized-open, 1.154 Å reduced-closed vs reduced-open, 1.424 Å oxidized-closed vs reduced-open). (II) Regardless of redox conditions or mutation, all closed conformations are globally nearly identical to each other, and the same applies to all of the open conformations (e.g. RMSDs: 0.523 Å oxidized-closed vs reduced-closed, 0.631 Å oxidized-open vs reduced-open, 0.808 Å reduced-open vs C283A,C284A-open). Notably, the distributions of dimers differ distinctly: for the WT under oxidizing conditions, 55% are closed and 41% open (4% too damaged to assign), whereas under reducing conditions, only 12% are closed and 88% are open (Table 2).

Observing the two different conformations in both WT datasets suggests that the enzyme samples the same global conformational equilibrium under both redox conditions. However, reducing conditions, which inhibit activity, heavily shift the equilibrium toward the globally open conformation. In contrast, under oxidizing conditions in which the enzyme is active, both global conformations appear to be sampled in nearly equal proportions. Further oxidation with diamide, beyond passive air oxidation, did not increase *Hn*CsoSCA’s specific activity (SI Fig. 1E), demonstrating that the WT oxidized sample used for cryo-EM and kinetics was already fully oxidized. The conformational ratios observed in the cryo-EM data therefore likely reflect the equilibrium distribution of a fully oxidized active enzyme. Moreover, the inactive C283A,C284A variant exists exclusively in the globally open conformation and thus appears unable to sample the closed conformation (Fig. 3E and F, SI Fig. 8C). Combined, these observations imply that adopting the globally closed conformation is required for catalysis and that conformational dynamics, specifically the ability to transition between the closed and open conformations, likely also play a critical role. They further suggest that redox conditions regulate this process by controlling the oxidation state of Cys283 and Cys284, with oxidizing conditions more readily enabling CsoSCA to adopt the closed conformation.

### CryoEM maps show signs of radiation damage

Based on the S-S distance and on the clear redox-dependent effect on both activity and conformation of WT *Hn*CsoSCA - both abolished by the C283A,C284A mutation - we expect the Cys283,Cys284 pair to form a disulfide bond under oxidizing conditions and to be reduced otherwise. Although the maps of the oxidized enzyme do not show density consistent with a disulfide bond (SI Fig. 9), this observation is likely explained by radiation damage during cryo-EM data collection, rather than by the absence of the disulfide itself. Disulfide breakage is common and among the earliest outcomes of beam-induced damage (36), having been reported at electron exposures lower than 10 e^-^/Å^2^ (37), i.e. far lower than the ∼65 e^-^/Å^2^ used to collect our data. Radiation damage has also been shown to occur more readily at redox-sensitive sites (38). Our data consistently follows this trend: for the non-regulatory Cys41,Cys49 pair, a disulfide is observed in the oxidized but not in the reduced structure (SI Fig. 10). As this non-regulatory disulfide appears less sensitive to radiation damage, it serves as an internal control of *Hn*CsoSCA’s overall redox state in the structures. Importantly, at cryogenic temperature in vitreous ice, radiation damage can cause local disulfide breakage but cannot trigger any large-scale conformational changes that would occur in solution when a disulfide is reduced. Thus, we expect the conformational populations observed for *Hn*CsoSCA in vitrified samples to reflect those present in solution, even if the Cys283,Cys284 disulfide is locally lost to radiation damage.

### The globally closed conformation arranges the active site for catalysis

To compare local features between the WT oxidized, the WT reduced and the mutant structures and thereby gain deeper insight into CsoSCA’s catalytic and regulatory mechanism, we superimposed all dimer structures using the Zn^2+^ coordination residues (Cys173, His242 and Cys253) as reference points (Fig. 5A). Within the active site, the three Zn^2+^-coordinating residues, the water molecule completing the Zn^2+^ tetrahedral coordination sphere, and the catalytic dyad (Asp175 - Arg177) align almost perfectly between all structures. Key differences include (Table 2): (I) His397 and Glu399 adopt a “near” position closer to the active site Zn^2+^ in the globally closed structures, whereas in the globally open structures both residues are shifted to a more distant “far” position. (II) In the oxidized-closed structure, His397 in the near position coordinates a water molecule, hereafter termed the active site water (W_AS_). No density for this water is observed in the reduced-closed structure or in any of the three globally open structures (oxidized-open, reduced-open and C283A,C284A-open). (III) The loop spanning residues 199-203 located just behind the active site adopts either a “closed” conformation, creating a tightly packed structure behind the Zn^2+^, or it is flipped in an “open” conformation. It is closed in the oxidized-closed and C283A,C284A-open structures, mixed (closed in one protomer, open in the other) in the oxidized-open structure, and open in both reduced structures.

**Figure 5:**
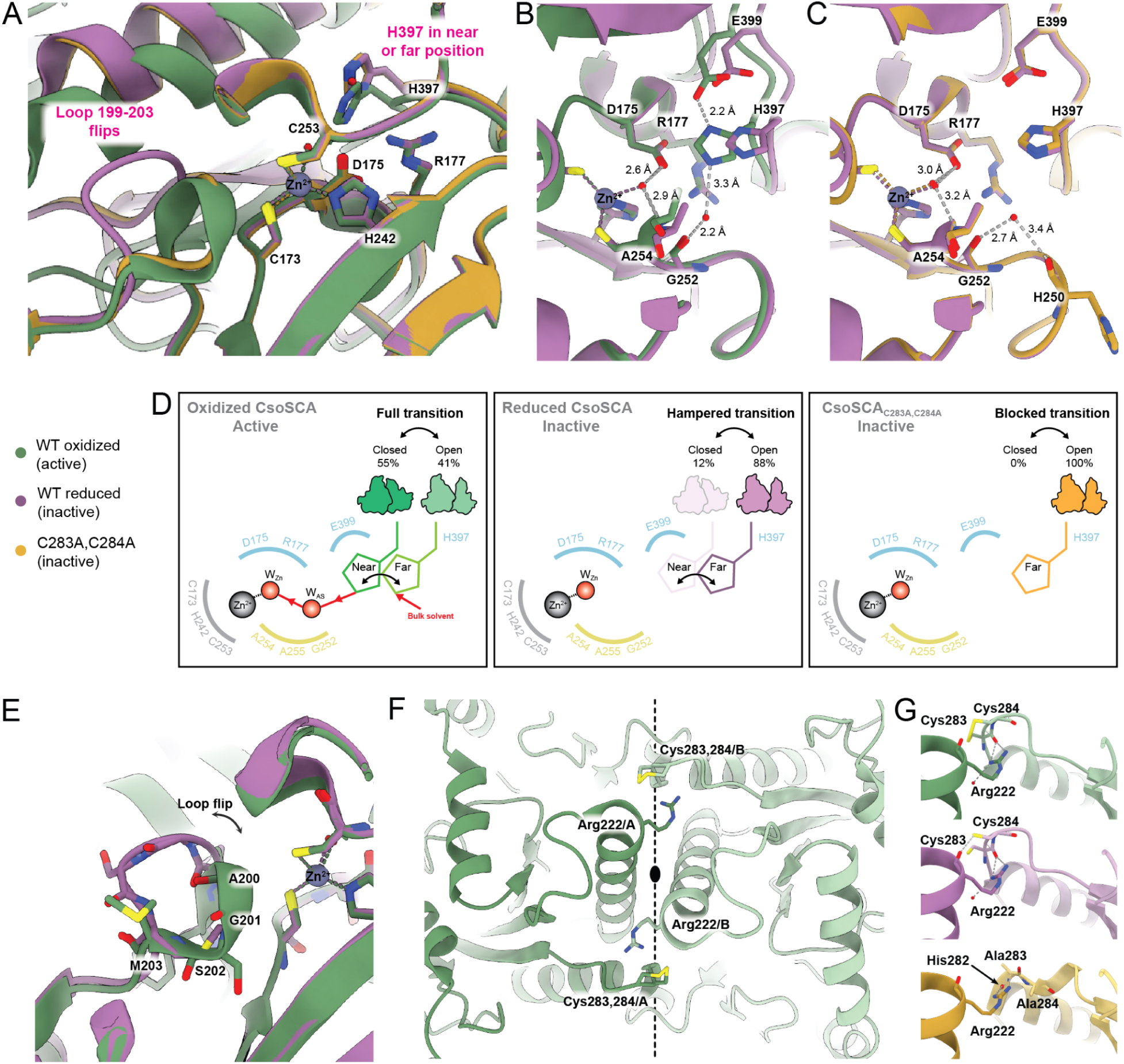
Structural details of the active site and regulatory cysteines explain the redox regulation. (A) Close-up view of the *Hn*CsoSCA active site and 199-203 loop. The WT oxidized-closed, WT reduced-open and C283A,C284A-open structures are superimposed based on the Zn^2+^-coordinating residues. Catalytically relevant residues are shown as sticks. (B-C) Close-up view of the active site comparing distances and the hydrogen bond network involved in catalysis: (B) oxidized-closed vs reduced-open structures and (C) reduced-open vs C283A,C284A-open structures. Hydrogen bonds are shown in grey and the indicated distances are those observed in the oxidized-closed (B) and reduced-open (C) structures. (D) Schematic of the *Hn*CsoSCA active site showing the near and far positions of His397, the observed water molecules and the catalytic Zn^2+^ for the three different structures. Arches indicate key active site residues, with hydrophilic residues in blue, hydrophobic residues in yellow and Zn^2+^-coordinating residues in grey. The proposed proton transfer pathway in the active oxidized condition is shown with red arrows. (E) Close-up view of the 199-203 loop in its closed conformation observed under oxidizing conditions (oxidized-closed structure) and its open conformation observed under reducing condition (reduced-open structure). Loop residues are shown as sticks. (F) View of the dimer interface showing the interaction between Arg222 from one protomer and the vicinal Cys283,Cys284 pair of the opposing protomer, in the oxidized-closed structure. Residues are shown as sticks and protomers are colored in dark (protomer A) and light (protomer B) green. The dimer’s 2-fold symmetry axis is indicated by a black ellipse and dashed line. (G) Close-up view of the interaction between Arg222 and the backbone at the vicinal Cys283,Cys284 pair in the oxidized-closed and reduced-open structures, and between Arg222 and the backbone at His282 in the C283A,C283A-open structure. Protomers are colored in dark and light shades of each structure’s color.

Rationalizing the regulatory mechanism based on these observations prompted us to first describe and update the catalytic mechanism of CsoSCA-catalyzed HCO_3_^-^ dehydration (Fig. 4A). As originally proposed by Sawaya et al (26), the catalytic cycle begins with Asp175 from the catalytic dyad displacing the Zn^2+^-coordinated water molecule (oxidized-closed, chain A: 2136). The substrate, HCO_3_^-^, then enters the active site and takes this place by coordinating to the Zn^2+^ and forming hydrogen bonds with Asp175, the backbone of Ala254 and Ala255, and Nε2 of His397, which position is stabilized by hydrogen-bonding to Glu399. This arrangement leads to decomposition of HCO_3_^-^ into CO_2_ and OH^-^. The product, CO_2_, diffuses out of the active site, likely via interactions with the Ala255 backbone within the hydrophobic side of the active site, consistent with a β-CA structure from *Pseudomonas aeruginosa* with bound CO_2_ (39). A water molecule then protonates the Zn^2+^-bound OH^-^, forming the product H_2_O and restoring the active site for the next cycle.

Previously, this proton-donating role was assigned to a water molecule hydrogen-bonded to Ala255 and proposed to interact with His397, but the 5.7 Å distance (water-His397) makes such an interaction unlikely. We therefore propose that the active site water observed in our oxidized-closed structure which hydrogen bonds to the backbone of Gly252 (2.2 Å) and to His397 (3.3 Å) mediates this proton transfer (Fig. 4B), with His397 functioning analogously to His64 in human CAII (a α-CA) by shuttling a proton from the bulk solvent into the active site (40, 41). All catalytically relevant residues are conserved (Fig 4C), and importantly, mutating His397 to alanine completely abolished catalytic activity (Fig. 4D, SI Fig. 1D), confirming its functional importance.

His397 and Glu399 in their near positions are essential for binding both HCO_3_^-^ and the active site water required for proton transfer. In our oxidized active structures, these positions occur only in the oxidized-closed conformation (Fig. 4B), and we therefore refer to it as the “catalytically ready” conformation. However, given that the oxidized dataset contains nearly equal proportions of globally open (far position His397/Glu399) and globally closed dimers (near position His397/Glu399) (Table 2), it is plausible that the movement of His397/Glu399 between these two positions is an essential step in the proton transfer, similar to the dynamics established for His64 in hCAII (41–45).

### The position of His397 is critical for catalysis and regulation

In light of the catalytic mechanism, the central role of His397 in positioning both HCO_3_^-^ and the active site water provides a key to understanding the redox-regulatory mechanism. As noted above, in the catalytically ready oxidized-closed structure, His397 in the near position coordinates the active site water (3.3 Å to Nε2), together with Gly252 (2.2 Å to backbone carbonyl) (Fig. 5A and B, Table 2). This near position of His397 is stabilized by hydrogen bonds between His397-Nδ2 and the Oε2 atom of Glu399 (2.2 Å). In contrast, in the inactive reduced-open and C283A,C284A-open conformations, both His397 and Glu399 occupy their far positions (Fig. 5A-C). At this position, the interaction between His397 and Glu399 is destabilized (3.4 Å reduced-open and 3.7 Å C283A,C284-open) and His397 is dislodged by more than 1 Å, placing it beyond hydrogen bonding distance to a potential active site water. In agreement with this, we observe no density for this water in the reduced-open structure. In the C283A,C284A-open structure, a water molecule is observed in one of the two protomers, but its distance to His397-Nε2 is increased to 4.2 Å which disrupts the His397-water-Gly252 hydrogen bonding network required for catalysis. Instead, the water is positioned further out from the active site via hydrogen bonds to the backbones of His250 and Gly252 (Fig. 5C). Although His397 is at its near position in the reduced-closed structure, no active site water density could be assigned in this structure.

In summary, this suggests a regulatory mechanism in which oxidation allows CsoSCA to fully transition between its open and closed conformations, allowing His397 to move between its far and near positions, enabling catalysis. Reduction or mutation of the regulatory cysteines hampers or blocks this transition, keeping His397 in the far position, preventing both HCO_3_^-^binding and proton transfer (Fig. 5D), thereby inactivating CsoSCA.

### The position of the 199-203 loop may be involved in catalysis and regulation

The other notable local difference is the position of the 199-203 loop, located just behind the active site Zn^2+^ (Fig. 5A and E, Table 2). This loop is composed mainly of hydrophobic residues, showing high degrees of conservation (Fig. 4C). Unlike His397 and Glu399, whose near and far positions correlate with the global conformation, the 199-203 loop is independent of this. In the catalytically ready oxidized-closed structure, it adopts a closed conformation with the C_α_ of the closest residue, Gly201, 4.9 Å from the Zn^2+^ (Fig. 5A and E). Notably, the oxidized-open structure shows a mixture of open/closed 199-203 loops. This indicates that either a closed loop, or its ability to switch between conformations during the catalytic cycle, may be critical for catalysis. In both reduced structures, the loop is flipped to an open conformation, increasing the Gly201-Zn^2+^ distance to 7.8 Å (Fig. 5A and E), suggesting that an open loop could be associated with an inactive state. However, the inactive C283A,C284A variant retains a closed loop (Fig. 5A), showing that the correct loop conformation alone is insufficient for activity.

### The regulatory vicinal cysteines control long range conformational dynamics

In the WT structures, the Cys283,Cys284 loop is stabilized by hydrogen bonds between the backbone carbonyl of Cys283 and the side chain of Arg222 from the other protomer (Fig. 5F and G). In the C283A,C284A variant, observed exclusively in the globally open conformation, this loop is more flexible, and Arg222 instead hydrogen bonds with the backbone carbonyl of His282 (Fig. 5G). This suggests that a more flexible 283-284 loop prevents the enzyme from sampling the closed conformation, resulting in an inactive state. Consistent with this, under reducing conditions the loop in the WT is expected to be more flexible than when a disulfide bond is present (46, 47). In this condition, our data show that *Hn*CsoSCA does not sufficiently sample the closed conformation and the enzyme is inactive, similar to the C283A,C284A variant, which is trapped in the open conformation. In contrast, under oxidizing conditions, when the enzyme is active, the 283-284 loop is likely more rigid, and *Hn*CsoSCA more readily adopts the closed conformation and samples both conformations at equal proportions.

## Discussion

In this study, we have elucidated the molecular mechanism underlying carbonic anhydrase regulation in α-carboxysomes. Through steady-state kinetic measurements, we found that the activity of the carboxysomal CA, CsoSCA, is regulated by the redox state of the environment: the enzyme is inactive under reducing conditions and active under oxidizing conditions. Bioinformatic analysis combined with targeted mutagenesis identified a conserved vicinal cysteine pair, located distal from the active site, as the key residues responsible for this regulation. Cryo-EM structures of WT CsoSCA in active and inactive conditions, along with an inactive cysteine mutant, reveal that redox conditions modulate global conformational changes that propagate to the active site. Together, these data explain how CsoSCA transitions from the inactive cytosolic state to the active state upon encapsulation in α-carboxysomes.

### Regulation of carboxysomal carbonic anhydrases

A requirement for the bacterial CCM to function is that CA activity is minimized in the cytosol and only located within carboxysomes (2, 24, 31–33). For the cell to accomplish this, the carboxysomal CA needs to be efficiently encapsulated and its activity tightly regulated. We previously discovered the encapsulation mechanism of the α-carboxysomal CA, CsoSCA, and that it occurs via multivalent interactions between its intrinsically disordered N-terminus and Rubisco (24). Here, we demonstrate that CsoSCA’s activity depends on its redox environment, revealing a mechanism for how α-carboxysomes accomplish this essential regulation.

Previous work using encapsulated redox-sensitive GFP has shown that carboxysomes maintain an internal oxidizing state, while the bacterial cytosol is kept in a reducing state due to the presence of cellular reductants (13–15). Oxidation of the interior is thought to occur through a maturation process after assembly in which the fully closed shell acts as a diffusion barrier, restricting entry of cellular reductants and resulting in gradual oxidation of the carboxysome lumen. In line with this model, our findings suggest a clear mechanism where CsoSCA is kept inactive in the reducing cytosol, preventing short-circuiting of the CCM, and only becomes active after encapsulation, when lumenal oxidation promotes disulfide formation. Hence, our data couple CsoSCA activity to carboxysome maturation and update the current model of α-carboxysomes assembly and activation as presented in Fig. 6.

Consistent with this, one of the two β-carboxysomal CAs, the γ-CA CcmM, is also redox-regulated (23). Notably, this shared regulatory mechanism occurs despite the fact that CsoSCA and CcmM are structurally and evolutionarily unrelated. As with CsoSCA, the inactivation of CcmM involves large redox-driven conformational changes that misposition catalytic residues and disrupt essential hydrogen bonding networks in the active site. However, how CcaA, the β-CA in β-carboxysomes (20, 22), and the recently identified α-carboxysome associated ι-CA (28) are regulated, and whether these are also redox-controlled, remains unknown.

The principle of redox-based control extends beyond CA activity, and redox-dependent changes have also been observed in protein-protein interactions required for the assembly of both α- and β-carboxysomes (12, 48–50), further underscoring that an oxidizing lumen is critical for proper carboxysome function. Taken together, these findings highlight that the effects on enzyme activity and protein interactions upon encapsulation within the specific microenvironment of bacterial microcompartments must be considered when studying and engineering carboxysomes and similar systems.

### Molecular mechanism of CsoSCA’s redox regulation

Based on our data, we propose that CsoSCA catalysis requires not only precise positioning of active site residues but also the ability to transition between the globally open and closed conformations. In the catalytically ready closed state, His397 and Glu399 adopt their near positions, enabling His397 to bind HCO_3_^-^ and, following bond breakage and CO_2_ release, coordinate the active site water that protonates the Zn^2+^-bound OH^-^. In the open state, both residues shift to their far positions, too far away to allow these interactions. Because the oxidized enzyme populates open and closed conformations in nearly equal proportions, we hypothesize that they are both essential for catalysis. In such a mechanism, His397 likely functions as a proton shuttle. In the far more solvent exposed position, His397 can acquire a proton from bulk solvent, which is then transferred to the active site water when the enzyme transitions back to the closed conformation (Fig. 4A and B, 5D). In α-CAs, a comparable functional role is well established for His64 in human CAII, which flips between solvent-exposed and inward-facing positions to mediate proton transfer (41–45). Fully establishing such dynamics and proton transfer mechanisms in CsoSCA will require similar extensive structural and biophysical characterization as has been performed for α-CAs.

Building on this mechanistic framework, we further propose that CsoSCA’s redox state controls this conformational dynamics and the ability to adopt the catalytically ready closed state. Under oxidizing conditions, formation of the Cys283,Cys284 disulfide at the dimer interface, ∼35 Å from the active site, allows sampling of both the open and closed conformations (Fig. 5D). Reduction of these cysteines strongly shifts the conformational equilibrium towards the open state, hampering His397 from adopting its near position and thereby abolishing activity. The C283A,C284A variant mirrors the reduced enzyme, remaining exclusively open and inactive, supporting such a mechanism. Together, these findings suggest that CsoSCA activity depends on its ability to access the closed conformation, which appears to be possible only when the enzyme is oxidized. The Cys283,Cys284 disulfide thus acts as a distal molecular switch, activating CsoSCA upon carboxysome encapsulation by controlling long-range conformational changes that shape the local active site architecture.

A direct mechanistic role for the 199-203 loop is difficult to define, as it is located behind the active site Zn^2+^ and does not directly contact catalytic residues. Nevertheless, its position follows the redox state - closed when oxidized and open when reduced - with some flexibility observed in the oxidized-open structure. However, Ala200 and Gly201 are part of defining the hydrophobic side of the active site proposed to form the CO_2_ binding pocket (39). In the reduced structures, opening of the loop alters this region, which could hinder catalysis. Notably, in all other available CsoSCA structures (26, 29, 30), as well as in canonical β-CAs (39, 51, 52), the loop consistently adopts the closed conformation. This may result from experimental conditions, or it may indicate that this conformation represents the active state.

Radiation damage, commonly affecting disulfides in both cryoEM and crystal structures (36–38, 53), most likely prevented us from visualizing the Cys283,Cys284 disulfide in our oxidized WT structure. A lower electron dose for data collection would likely not have helped to reveal it, as sensitive disulfides can be damaged before 10 e/Å^2^ (37, 38), a dose too low to provide sufficient contrast for particle picking. However, the more resistant Cys41,Cys49 disulfide is resolved under oxidizing conditions, verifying the redox state in the structures. In the future, zero-dose extrapolation (54) has the potential to visualize radiation-sensitive features, and its implementation in image processing programs will provide a new tool to study such fragile regulatory disulfides.

### Evolutionary differences within carboxysomal CAs

The catalytic constants determined for *Hn*CsoSCA and *Cy*CsoSCA fall within the same range, indicating that this activity level is typical for α-carboxysomal CAs. Interestingly, the *k*_cat_ of ∼10^4^ s^-1^ is similar to that of β-carboxysomal CAs (19, 20), suggesting that this rate - modest compared to many other CAs - represents the activity necessary to sustain the bacterial CCM. Further, our kinetic and bioinformatic analysis shows that redox regulation is a conserved feature of CsoSCA, present in all cyanobacterial and most proteobacterial homologs. However, three distinct evolutionary clades (Fig. 2E, SI Fig. 3) lack the redox-switching cysteines. It remains to be determined whether these enzymes retain redox sensitivity through alternative residues, employ a different regulatory mechanism, or simply do not require cytosolic inactivation of CA under their environmental conditions. Redox regulation has also been described in other CAs, including variants from plants, algae and diatoms: however, in some of these enzymes the relationship is reversed, with oxidation causing inactivation and reduction restoring activity (55–58).

In addition to redox control, cyanobacterial CsoSCA features a second regulatory layer: allosteric activation by the Rubisco substrate RuBP, proposed to fine-tune the internal pH of carboxysomes during fluctuating RuBP concentrations in cyanobacteria (29, 59). This allosteric site is located in the C-terminal domain. However, as no structure of apo-*Cy*CsoSCA is available, the molecular basis for this regulation, and whether it involves movement of the same active site residues (His457 and Glu459 in *Cy*CsoSCA, corresponding to His397 and Glu399 in *Hn*CsoSCA’s) that mediate the redox control, remains unresolved. By contrast, the function of the C-terminal domain in proteobacterial CsoSCA is unknown, as is whether these organisms require internal carboxysome pH regulation through a mechanism analogous to that in cyanobacteria.

In summary, this work advances our understanding of carboxysome regulation focusing on the essential CA and the functional importance of oxidation of the mature carboxysome. These insights are critical for efforts to engineer carboxysome-based CCMs into non-native hosts to improve growth and productivity in crops and industrially important microorganisms (60, 61). More broadly, we hope that these findings will deepen our mechanistic understanding of enzyme regulation and how catalytic activities are affected upon encapsulation into different types of compartments, thereby supporting the development of future biotechnological applications.

## Materials and Methods

### Bioinformatics

The 222 CsoSCA protein sequences collected in Blikstad et al 2023 (24) were re-analyzed to identify potential regulatory cysteines, with the minor difference that the correct full-length *Cy*CsoSCA was used after its previously misannotated start codon was clarified in (29). Sequences were first aligned using MUSCLE (62). The resulting MSA was used to build a phylogenetic tree using IQ-TREE web server (63) and visualized using iTOL (64). Sequence logos of the MSAs were visualized with WebLogo3 (65). Conserved motifs in the unstructured NTD of CsoSCA were analyzed with The MEME suite (66).

### Protein expression and purification

All CsoSCA constructs were recombinantly expressed as N-terminal Strep-MBP fusions using a pET14 expression plasmid transformed into *E. coli* BL21-AI cells. For protein expression, cells were grown in LB-medium supplemented with 100 μg/mL ampicillin at 37 °C. At OD_600_ = 0.6 - 0.8 the expression was induced with addition of 0.1 % L-arabinose and supplemented with 100 μM ZnSO_4_ to ensure full incorporation of Zn^2+^ in the active site. The temperature was decreased to 18 °C, cells were grown overnight and subsequently harvested by centrifugation at 18,500 x *g* (Rotor JA-10, Avanti J-25 Beckman Coulter). Bacterial pellets were frozen at −20 °C until use.

All proteins were purified with Strep-tag affinity chromatography, with all steps carried out at 4°C. Bacterial pellets were thawed and resuspended in purification buffer (50 mM Tris, 300 mM NaCl pH 8.0), supplemented with 0.1 mg/mL lysozyme, 0.1 μg/mL benzonase (Merck) and 1 mM phenylmethylsulfonyl fluoride (PMSF) and lysed by sonication at 65% amplitude with two rounds of 10 s pulse 15 s pause for 1 min (VibraCell, Sonics). Lysed cells were clarified by centrifugation at 27,670 x *g* for 45 min, at 4 °C. The clarified lysates were then applied to a 5 mL StrepTrap HP column (Cytiva) pre-equilibrated with purification buffer using a syringe pump (AL-4000, KF technologies) at a flow rate of 2.5 mL/min. Unspecific proteins were washed away with 60 mL of purification buffer at a flow rate of 5 mL/min, and the bound proteins were thereafter eluted with 3 x 10 mL purification buffer supplemented with 2.5 mM desthiobiotin at a flow rate of 2.5 mL/min. The pure protein samples were thereafter concentrated by ultrafiltration to 2.5 mL using a 30 kDa cut-off filter (Amicon Ultra-4 30,000 MWCO, Milllipore) and buffer exchanged into storage buffer (50 mM Tris, 150 mM NaCl, pH 7.5), using a PD-10 column (Cytiva). Final purified proteins were stored at 4 °C until further use. It should be noted that addition of glycerol and storing the samples at −80 °C negatively impacts the activity of the enzyme. Similarly, storage at 4 °C was limited to 7 days post-purification before loss in activity was observed. All kinetic measurements were therefore performed within 5 days of purification and vitrification for structure determination was done within 2 days. Purity was checked with SDS-PAGE and was in general >95% pure. Protein concentrations were determined spectrophotometrically at A_280_ using a NanoDrop2000 (Thermo Scientific) and the theoretically calculated extinction coefficients (Expasy ProtParam, *Hn*CsoSCA ε: 120,140 M^-1^ cm^-1^ and *Cy*CsoSCA ε: 115,210 M^-1^ cm^-1^).

### Carbonic anhydrase kinetic measurements

#### Saturation curve

Steady-state kinetic parameters for *Hn*CsoSCA and *Cy*CsoSCA were determined by monitoring the hydration of CO_2_ using the Khalifah/pH indicator assay (34), using a SX20 stopped-flow spectrophotometer (Applied Photophysics). For a detailed protocol see Vogiatzi and Blikstad, 2024 (35). Briefly, in the experimental set-up: syringe 1 containing assay buffer, pH indicator and enzyme and syringe 2 containing CO_2_, were mixed in a 1:1 ratio. All measurements were conducted at 25 °C. Measurements were performed at pH 8.0 using the buffer-indicator pair HEPES-phenol red monitored at 558 nm. Final buffer concentration was 50 mM, with the total ionic strength adjusted to 50 mM using Na_2_SO_4_, and final indicator concentration at 50 μM. For *Hn*CsoSCA the final enzyme concentration was 84 nM (monomer) and for *Cy*CsoSCA 88 nM (monomer). For *Cy*CsoSCA, measurements were performed with addition of 150 μM RuBP. The final concentration of CO_2_ was 1.7 - 17 mM. A saturated solution of CO_2_ (34 mM at 25 °C) was prepared by continuously bubbling CO_2_ into milli-Q water at 25 °C. To obtain lower concentrations the saturated CO_2_ solution was diluted in milli-Q water using a 2.5 mL gas-tight Hamilton syringe. Initial rates (0 - 0.5 s) were measured in 8 replicates, care was taken to keep the change in absorbance of initial rates below 10% of the total reaction. The total reactions until equilibrium (0 - 60 s) were measured in duplicates and used to calculate experimentally determined buffer factors (linear relationship between total absorbance change vs total change in CO_2_ concentration) for converting the change in absorbance to the change in CO_2_ concentration. To extract the kinetic constants *k*_cat,_ *K*_M_ and *k*_cat_/*K*_M_, the Michaelis-Menten equation was fitted to background-subtracted initial rates using non-linear regression using MMFIT (for *k*_cat_ and *K*_M_) and RFFIT (for *k*_cat_/*K*_M_) in the SimFIT package (https://simfit.uk).

#### Specific activity measurements to test redox dependence and mutants

To test redox dependence and influence of site-specific mutations, specific activity was measured at 17 mM CO_2_, using the method described above. In all cases, measurements were performed at 25 °C in pH 7.5 using the buffer-indicator pair MOPS-para-nitrophenol (pNP) monitored at 400 nm. Final buffer concentration was 50 mM, with the total ionic strength adjusted to 50 mM using Na_2_SO_4_, and final indicator concentration at 50 μM. Measurements in oxidizing conditions were performed in the aforementioned assay buffer without supplement. For the reducing conditions, the assay buffer was supplemented with 1 mM tris-(2-carboxyethyl)phosphine (TCEP) and the samples were incubated for 15 min prior to measurement. For further oxidation of *Hn*CsoSCA, the assay buffer was supplemented with 1 mM of diamide (tetramethylazodicarboxamide) and the samples were incubated 15 min prior to measurement. For site-specific mutations, the following cysteine mutants and active site histidine mutant were tested: *Hn*CsoSCA_C283A_, *Hn*CsoSCA_C284A_, *Hn*CsoSCA_C283A,C284A_, *Hn*CsoSCA_C41A_, *Hn*CsoSCA_C49A_, *Hn*CsoSCA_H397A_ and *Cy*CsoSCA_C353A,C354A_. All measurements were performed at an enzyme concentration of 0.5 μM (monomer) and in 3 technical replicates, where one replicate consists of an average of 8 traces for initial rates (0.0 - 0.5 s) and two traces for total reaction (0.0 - 60 s). To confirm loss of biologically relevant activity of the WT reduced sample and the mutants all measurements were repeated with 10-fold higher (5 μM) enzyme concentration.

### Cryo-EM specimen preparation and screening

Quantifoil R 1.2/1.3 300 Cu grids were glow-discharged in a Pelco easiGlow (20 mA, 60 s, 0.4 mbar of residual ambient air, negative polarity). For all conditions, 4 μL of protein solution was applied to the carbon side of the grid before blotting. All proteins came from the same preparations used for activity measurements, and were vitrified on the same day their activity was measured. MBP-*Hn*CsoSCA in oxidizing condition (no supplement) was at 0.7 μM (hexamer). MBP-*Hn*CsoSCA in reducing conditions was at 1.3 μM (hexamer) and supplemented with TCEP to a final concentration of 1 mM. The reduced sample was incubated 15 min with TCEP prior to vitrification. MBP-*Hn*CsoSCA_C283A,C284A_ was at 0.7 μM (hexamer) in oxidizing condition (no supplement). The blotting and plunge-freezing were performed with a Vitrobot Mark IV device, with the chamber at 4 °C and 100% relative humidity, no delay time (incubation between application and blotting), and 3 s of blotting (at blot force 0) before vitrification in liquid ethane. Multiple grids for each condition were screened on a Glacios microscope, operated at 200 kV with a Falcon3 camera operated in linear mode, to identify a suitable grid for high-resolution data collection.

### Cryo-EM data collection

The data were collected on a Titan Krios G2 operated at 300 kV and equipped with a Gatan K3 BioQuantum detector operated in counting mode. The microscope was operated with EPU version 3.6.0. The nominal magnification of 130 000x resulted in a raw pixel size of 0.65 Å/px. The energy filter slit width was 20 eV. The nominal defocus range was −1.8 to −0.6 μm by steps of 0.2 μm. The total dose was 64.76 e^-^/Å^2^ for the active and reduced *Hn*CsoSCA, 68.90 e^-^/Å^2^ for *Hn*CsoSCA_C283A,C284A_, and fractionated into 40 movie frames for all three datasets. Detailed data collection parameters for all three datasets are listed in Table S1.

### Cryo-EM image processing

Image processing statistics for all structures are listed in Table S1.

#### Pre-processing, particle picking and globally averaged 3D refinement

Image processing was performed with CryoSPARC version 4.7.0 (67). The same strategy was applied to all three datasets. A summary of the processing strategy is shown in SI Fig. 4-6. Raw movies were motion corrected using the Patch Motion Correction job with default parameters. CTF parameters were estimated using the Patch CTF estimation job with default parameters. The resulting micrographs were grouped based on the aberration-free image shift (AFIS) groups from data collection. Micrographs with a CTF fit resolution worse than 5 Å were excluded from further processing. About 1000 particles were manually picked to produce a training set for Topaz. A picking model was trained using the ResNet8 architecture, a micrograph downscaling factor automatically calculated by the CryoSPARC wrapper from the raw pixel size and a particle diameter of 150 Å, and an expected number of particles per field of view of 200, using topaz version 0.2.5 (68). Picked particles were extracted in a box 480 px wide, Fourier cropped to 72 px (resulting in 4.33 Å/px), and subjected to reference-free 2D classification (100 classes, circular mask diameter 160 Å). False positives from picking were excluded, and true particles retained, based on the respective absence or presence of recognizable protein features in their 2D class averages. These particles were re-extracted in a box 480 px wide Fourier cropped to 128 px (resulting in 2.44 Å/px), and sorted into 4 classes by *ab initio* reconstruction followed by heterogeneous refinement. Particles from the good classes were re-extracted in a box 480 px wide at the original pixel size (0.65 Å/px) and subjected to homogeneous refinement with D3 symmetry and optimization of per-particle scales, per-particle defocus and per-group CTF parameters (tilt/trefoil) (69). The reconstruction at this stage reached a resolution better than 3 Å for all datasets, and was used to perform reference-based motion correction (70) of the entire particle stack. The resulting motion-corrected particles were subjected to non-uniform refinement (71) with D3 symmetry and optimization of per-particle scales, per-particle defocus and per-group CTF parameters (tilt/trefoil), resulting in the hexamer reconstruction. Inspection of this reconstruction identified sharp features at the trimerization interface, but significant blurring at the outer surface of the protein. In addition to these observations, 3D variability analysis (72) solving for 3 principal components of variability with a filter resolution of 5 Å revealed significant motion of the hexamer (Movie 1, 2, 3), likely explaining the observed blurring.

**Figure 6:**
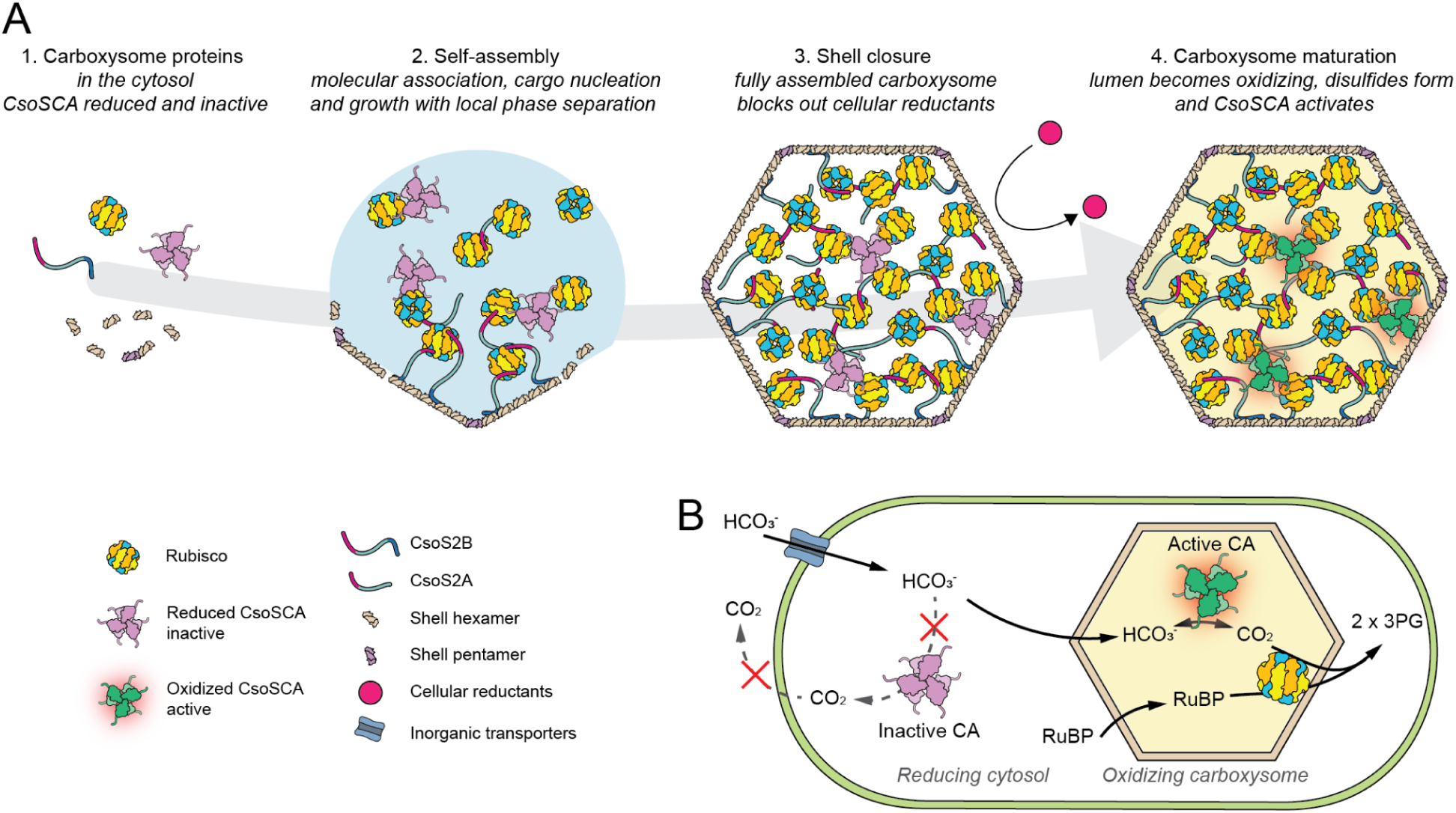
Updated model of the α-carboxysome based CCM. (A) Schematic of α-carboxysome assembly followed by oxidation of the lumen and activation of CsoSCA. The model includes: 1) CsoSCA being reduced and inactive in the cytosol. 2) Self-assembly through molecular association of carboxysome proteins, cargo nucleation and growth with local phase separation. 3) Shell closure forming fully assembled carboxysomes. The closed shell excludes cellular reductants. 4) The lumen of the carboxysome oxidizes, disulfides form and CsoSCA activates, resulting in mature and active carboxysomes. B) Schematic model of the α-carboxysome based CCM. The reducing cytosol keeps the CA inactive in the cytosol and prevents short-circuit of the CCM. The CA localized inside the oxidizing carboxysome is active and equilibrates HCO_3_ ^-^ to CO_2_ , thereby supplying Rubisco with a high concentration of CO.

#### Classification of dimers with different conformations

Initial atomic model building and refinement in the globally averaged, D3-symmetric hexamer map indicated that large regions of the map support a mixture of conformations. From this initial atomic model, we generated a mask encompassing a single dimer. The mask was generated using the molmap command from ChimeraX (73) with a resolution of 15 Å, imported into CryoSPARC and processed (binarized with a threshold of 0.02, dilated with a radius of 3 px, soft padded with a width of 14 to 18 px depending on resolution of the corresponding map following the equation from the CryoSPARC documentation). The set of particles from the globally averaged, D3-symmetric reconstruction was expanded in the lower order C3 symmetry to define the dimer as the asymmetric unit and generate particle metadata corresponding to all dimers. These dimer subparticles were then subjected to masked 3D classification with no alignment (under the assumption that the global alignments are accurate, a safe assumption given the high resolution of the D3-symmetric reconstruction) into 10 classes, using the mask defined above, with auto-tuning of class similarity. This procedure sorted all dimers based on their conformational heterogeneity. The resulting classes revealed some dimers visibly denatured (with missing density), likely by collisions to the air-water interface during blotting (74). Populations of dimers with different global conformations were also identified. Particles from each class were subjected to signal subtraction and local refinement (using a pose/shift gaussian prior during alignment, with default settings for the standard deviations of the shift and rotation priors, and default search ranges for local shift and rotation searches). The resulting locally refined dimer reconstructions were compared and regrouped into two classes based on their similarity: one major class globally similar in conformation to the consensus dimer, and one minor class markedly different from the consensus. For MBP-*Hn*CsoSCA_C283A,C284A_, 3D classification only revealed a single global conformation resembling the major class of the reduced WT enzyme.

#### Local resolution estimation

Local resolution values were estimated using a local FSC threshold of 0.143 in CryoSPARC, and are reported as min/median/max values inside the mask used in 3D refinement.

##### Atomic model building, refinement and validation

An initial model was prepared using AlphaFold2-multimer (75) to generate a prediction of the *Hn*CsoSCA dimer. This initial atomic model was rigidly placed into the map with ChimeraX version 1.10 (73), and the disordered N-terminus segment not supported by density was trimmed. The model was then fitted to the map by interactive molecular dynamics flexible fitting (iMDFF) using ISOLDE version 1.10 (76). Every residue in the *Hn*CsoSCA dimer was visually inspected at least once. The model was finally refined against the half-maps using Servalcat version 0.4.99 (77) (with riding hydrogens and isotropic B-factors). The same model building and refinement strategy was followed for all dimer maps. The hexamer maps were modeled by rigidly placing the dimer model of the most closely matching conformation, followed by refinement with C3 symmetry by Servalcat. Model validation and model-to-map fit analysis were performed with phenix.molprobity and phenix.validation_cryoem from the Phenix suite version 1.21.2-5419 (78, 79). In the oxidized-closed structure, a disulfide between Cys283 and Cys284 was automatically annotated by the PDB based on the S-S distance. Despite no support in the map (which we attribute to radiation damage, see SI Fig. 9 and 10), we chose to leave this annotation because it is consistent with the observed redox-dependent activity. Model building and refinement statistics for all structures are listed in Table S1.

##### Visualization and structural analysis

Structural figures were prepared using ChimeraX version 1.10 (73).

## Data availability

Cryo-EM maps and masks have been deposited in the Electron Microscopy Data Bank, and the corresponding atomic models have been deposited in the Protein Data Bank. Accession codes: *Hn*CsoSCA-WT in oxidizing conditions, hexamer (pdb_00009skr, EMD-54972), dimer major class (pdb_00009sks, EMD-54973), dimer minor class (pdb_00009skt, EMD-54974); *Hn*CsoSCA in reducing conditions, hexamer (pdb_00009sku, EMD-54975), dimer major class (pdb_00009skv, EMD-54976), dimer minor class (pdb_00009skw, EMD-54977); *Hn*CsoSCA-C283A-C284A, hexamer (pdb_00009skx, EMD-54978), dimer state 1 (pdb_00009sky, EMD-54979), dimer state 2 (pdb_00009skz, EMD-54980). Raw cryo-EM datasets were deposited to EMPIAR: *Hn*CsoSCA in oxidizing conditions (EMPIAR-12991), *Hn*CsoSCA in reducing conditions (EMPIAR-12992), *Hn*CsoSCA-C283A-C284A (EMPIAR-12993). Plasmids for all protein constructs used are available upon request. All protein sequences used for bioinformatics are available in FASTA format in Dataset S1.

## Acknowledgments

We are grateful to Karin Stensjö for helpful discussions and to Mikael Widersten for advice and access to the stopped-flow instrument. We acknowledge the use of the Cryo-EM Uppsala facility for specimen vitrification, grid screening, data storage and computing, funded by the Department of Cell and Molecular Biology, the Disciplinary Domains of Science and Technology and of Medicine and Pharmacy at Uppsala University. The data were collected at the Cryo-EM Swedish National Facility funded by the Knut and Alice Wallenberg, Family Erling Persson and Kempe Foundations, SciLifeLab, Stockholm University and Umeå University. We thank Dustin R. Morado and Julian Conrad for assistance with cryo-EM data collection. This work was supported by grants from the Lars Hierta Memorial Foundation, the O. E. and Edla Johansson Scientific Foundation and the Helge Ax:son Johnson Foundation to GG. This work was funded by grants from the Swedish Research council (2019-03700 and 2023-05296) and FORMAS - the Swedish Research Council for Sustainable Development (2019–01171) to CB.

## Author contributions

NV and CB designed research. NV, JL, TA and TS performed biochemistry. NV and GG performed cryo-EM and model building. NV, GG, JL, TA, TS and CB analyzed data. NV, GG and CB wrote the paper with input from all co-authors. GG and CB obtained funding. CB provided supervision.

## Supplemental Information

**SI Figure 1:**
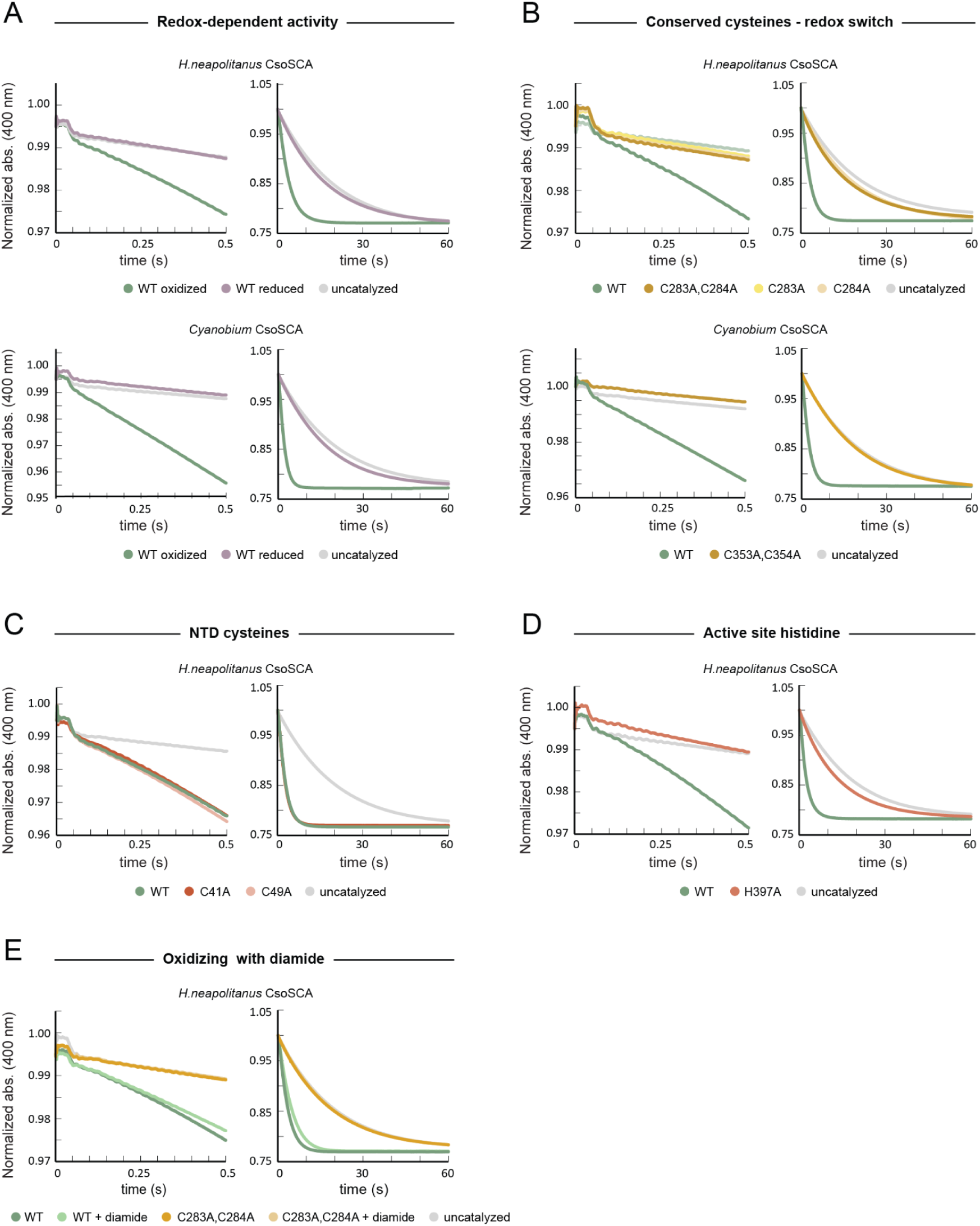
Specific activity measurements using 0.5 μM (monomer) of *Hn*CsoSCA and *Cy*CsoSCA. (A) Activity in oxidizing vs reducing conditions, showing that both *Hn*CSoSCA and *Cy*CsoSCA are only active in an oxidizing environment. (B) Alanine substitutions of the conserved cysteine pair abolish activity in both enzymes, showing that this cysteine pair acts as a redox switch. (C) Alanine substitutions of the NTD cysteines in *Hn*CsoSCA retain WT-like activity. (D) The active site variant H397A completely abolishes activity in *Hn*CsoSCA. (E) Additional oxidation with diamide does not increase WT activity of *Hn*CsoSCA or rescue the activity of the C283A,C284A variant. All measurements were performed in triplicates, using the MOPS-pNP buffer-indicator pair at pH 7.5. Left panels show the short traces (0.5 s), measuring initial rates at steady-state. Right panels show the long traces (60s), measuring full reaction until equilibrium.

**SI Figure 2:**
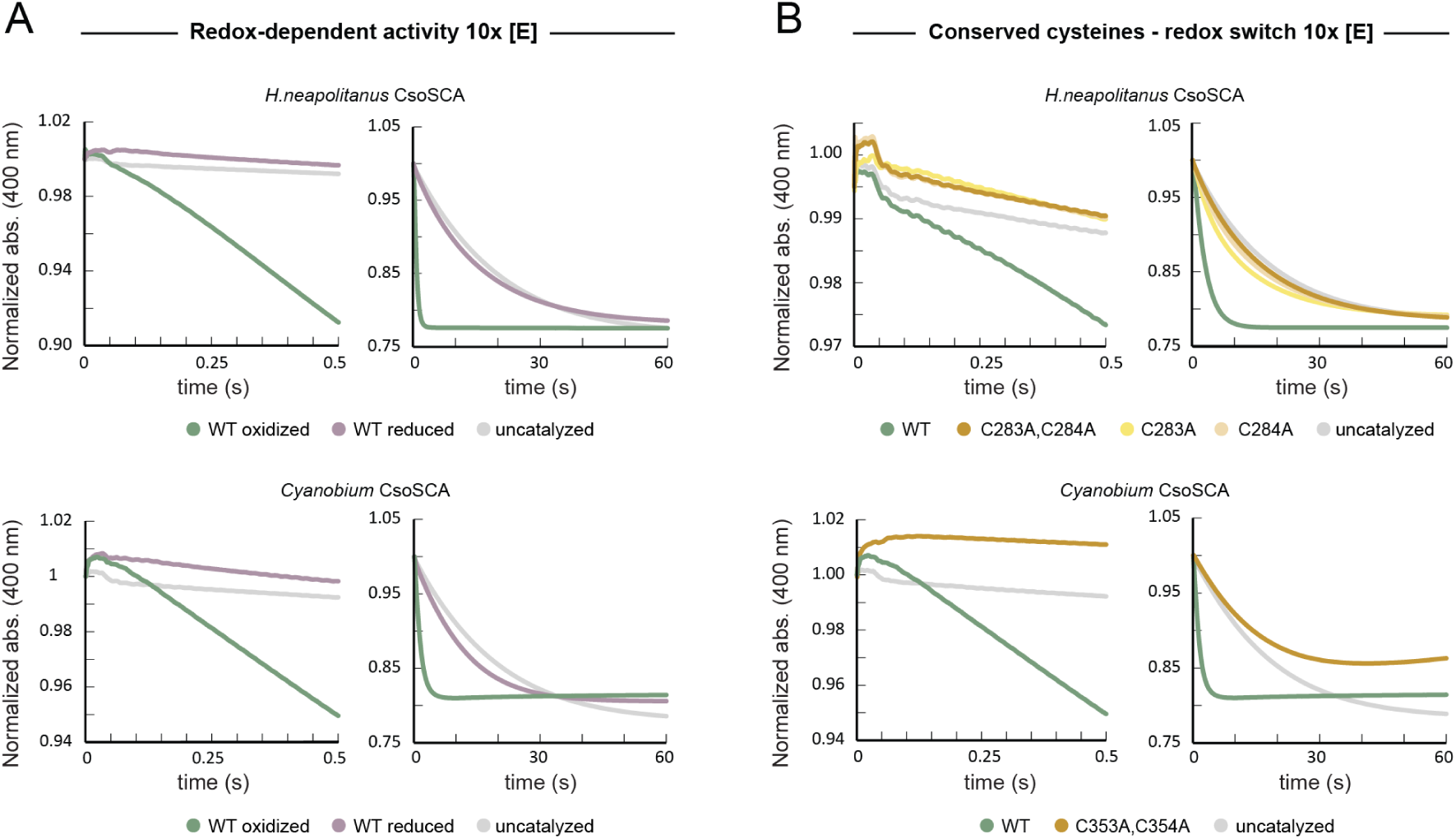
Specific activity measurements using 10x [E] (5 μM monomer) of *Hn*CsoSCA and *Cy*CsoSCA. (A) Activity in oxidizing vs reducing conditions using 10 times higher enzyme concentration than used in SI Fig 1A, verifying the absence of physiologically relevant activity in reducing conditions. (B) Activity of alanine substitutions of the conserved cysteine pair using 10 times higher enzyme concentration than used in SI Fig 1B, verifying that these variants are inactive. All measurements were performed in triplicates, using the MOPS-pNP buffer-indicator pair at pH 7.5. Left panels show the short traces (0.5 s), measuring initial rates at steady-state. Right panels show the long traces (60s), measuring full reaction until equilibrium.

**SI Figure 3:**
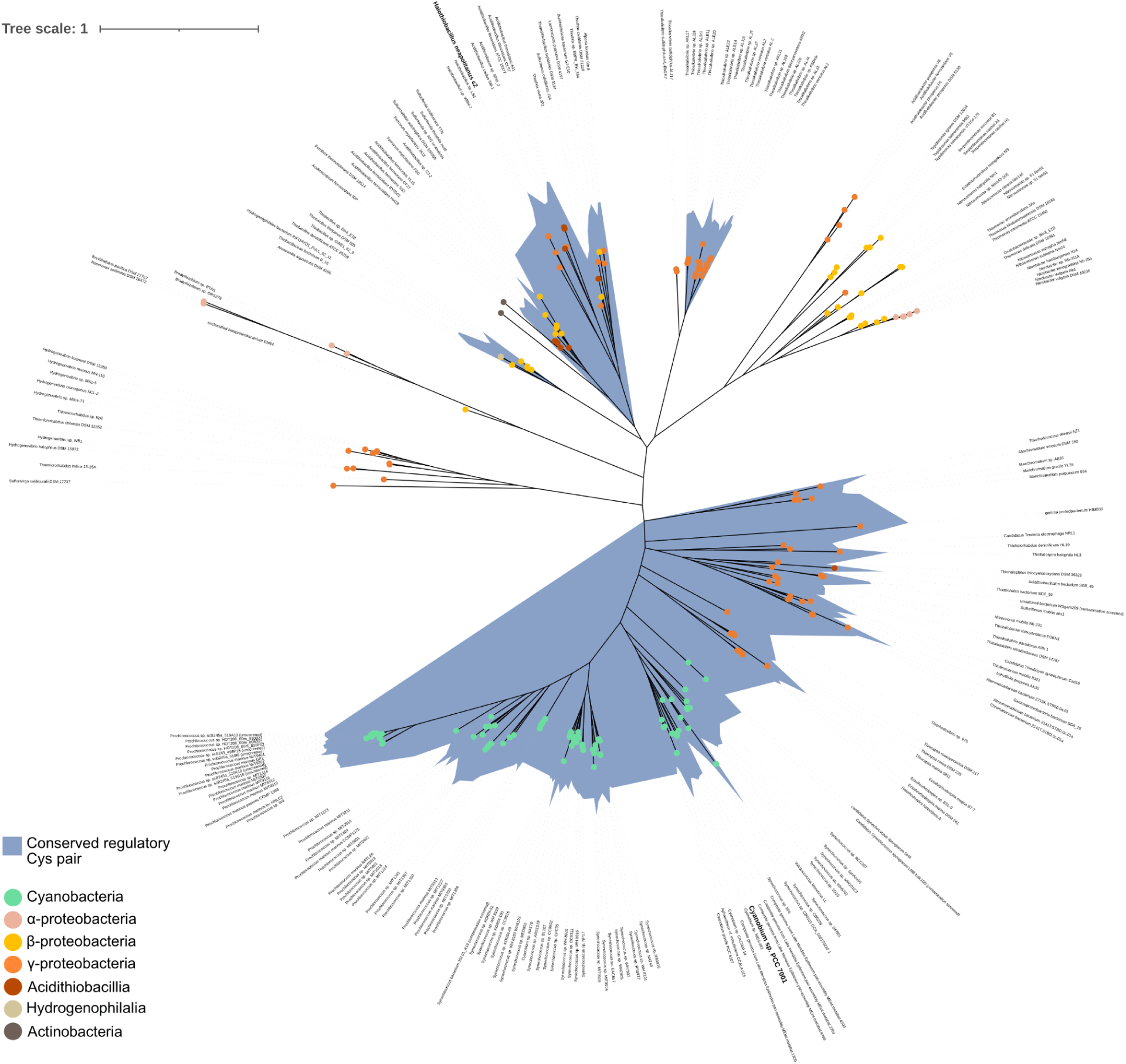
Annotated unrooted maximum-likelihood phylogenetic tree of CsoSCA. Clades containing the conserved regulatory cysteine pair are highlighted in blue. Scale bar 1 substitutions per site.

**SI Figure 4:**
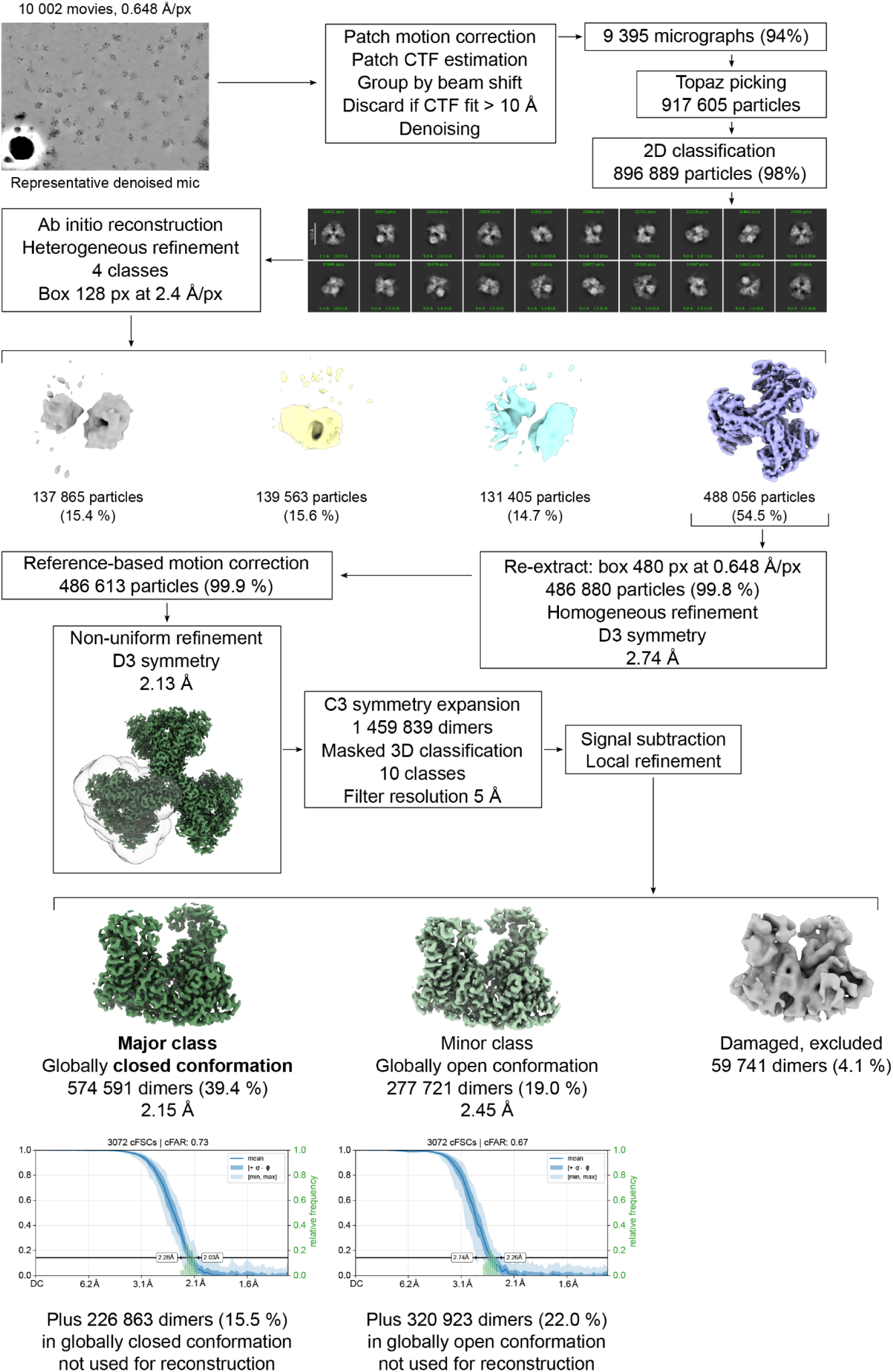
Cryo-EM image processing of *Hn*CsoSCA in oxidizing conditions.

**SI Figure 5:**
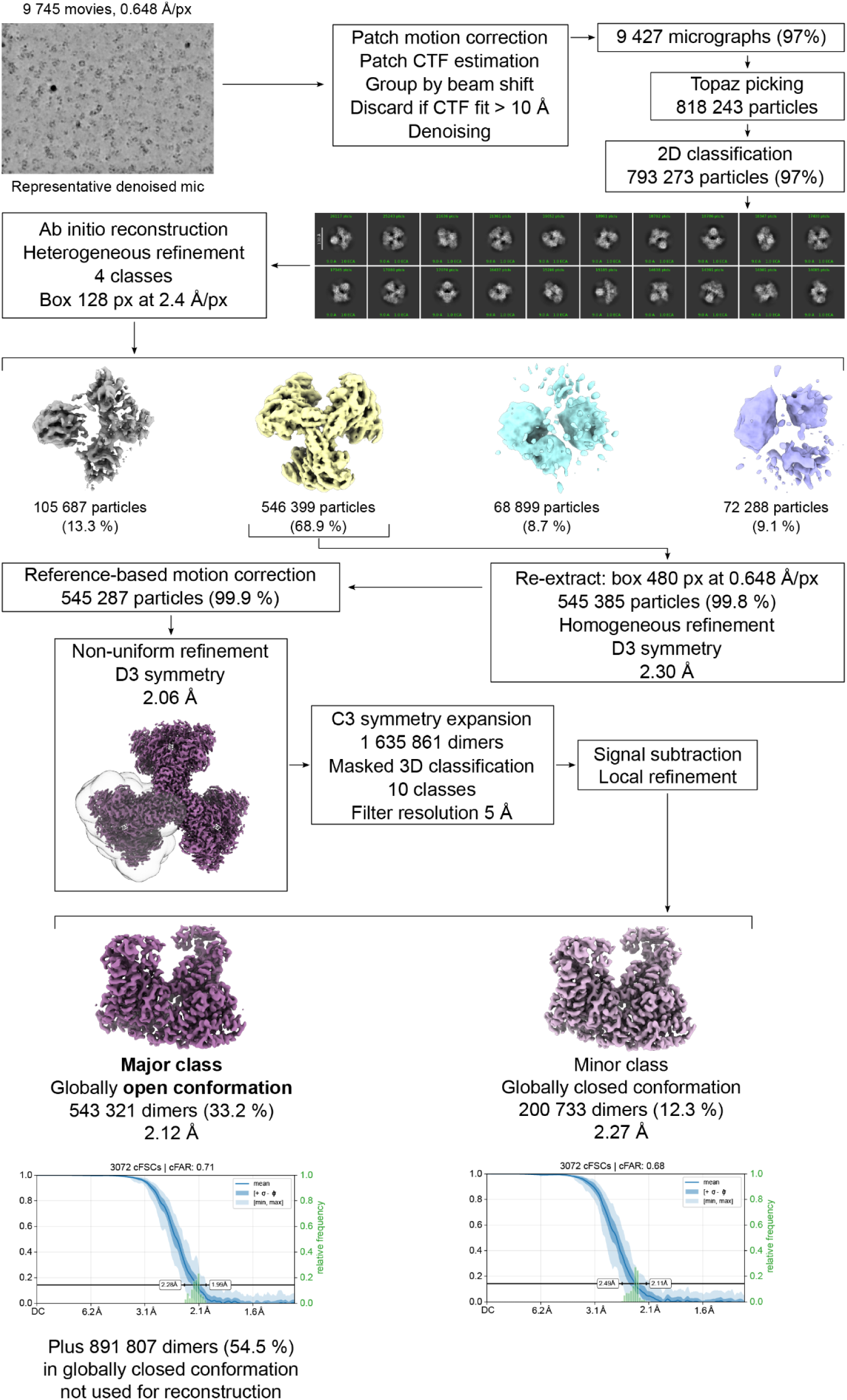
Cryo-EM image processing of *Hn*CsoSCA in reducing conditions.

**SI Figure 6:**
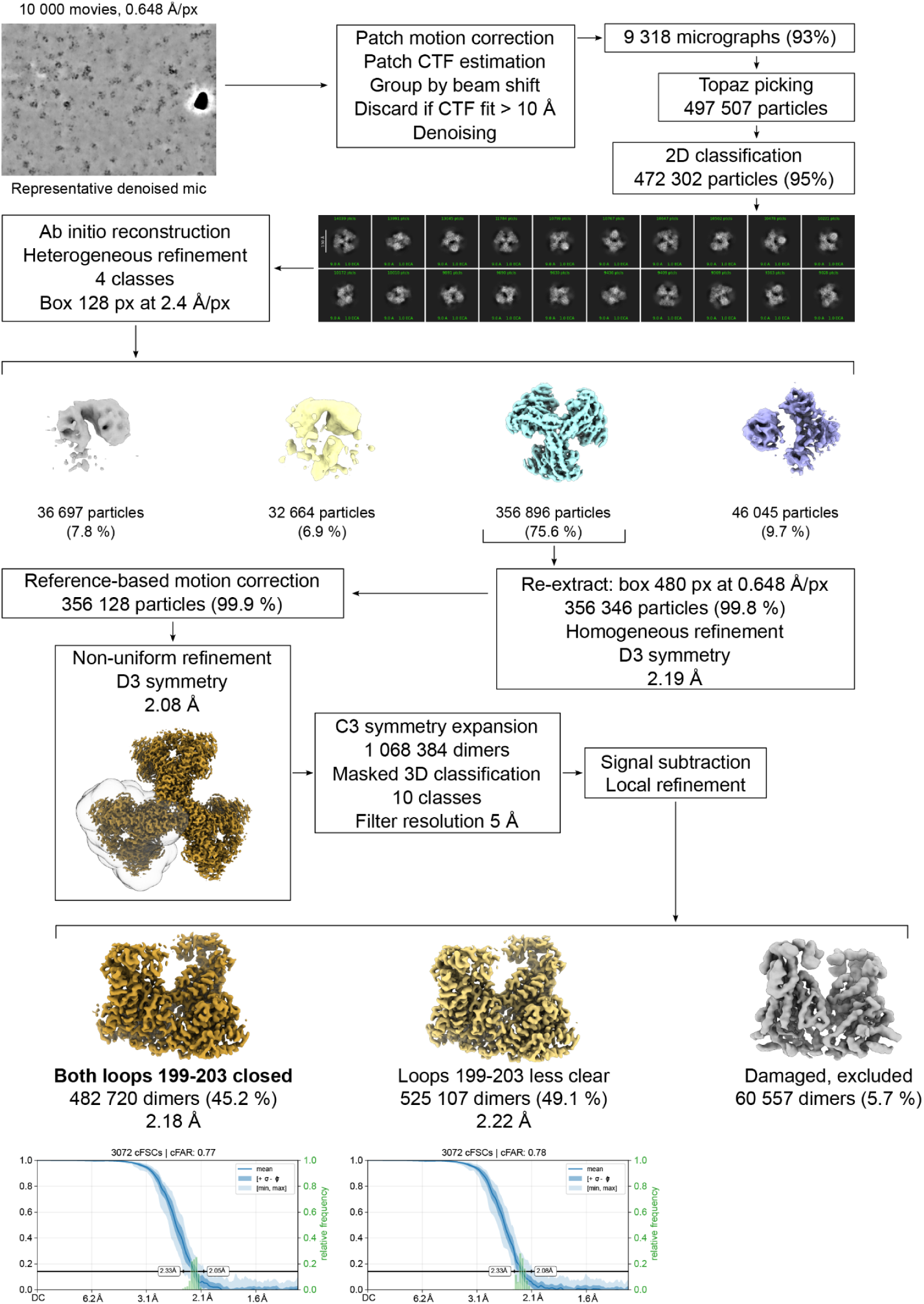
Cryo-EM image processing of *Hn*CsoSCA_C283A,C284A_ in oxidizing conditions.

**SI Figure 7:**
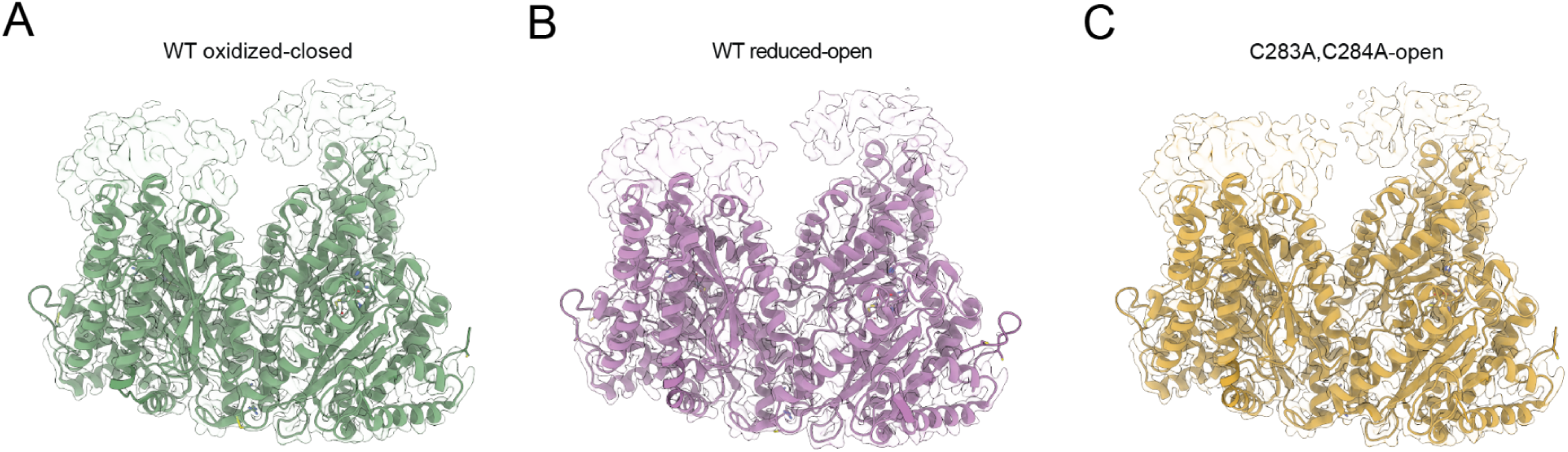
Atomic models of the *Hn*CsoSCA dimer structures. For each model, the corresponding cryo-EM map is overlaid and shown as a translucent surface. (A) Oxidized-closed structure. (B) Reduced-open structure. (C) C283A,C284A-open structure.

**SI Figure 8:**
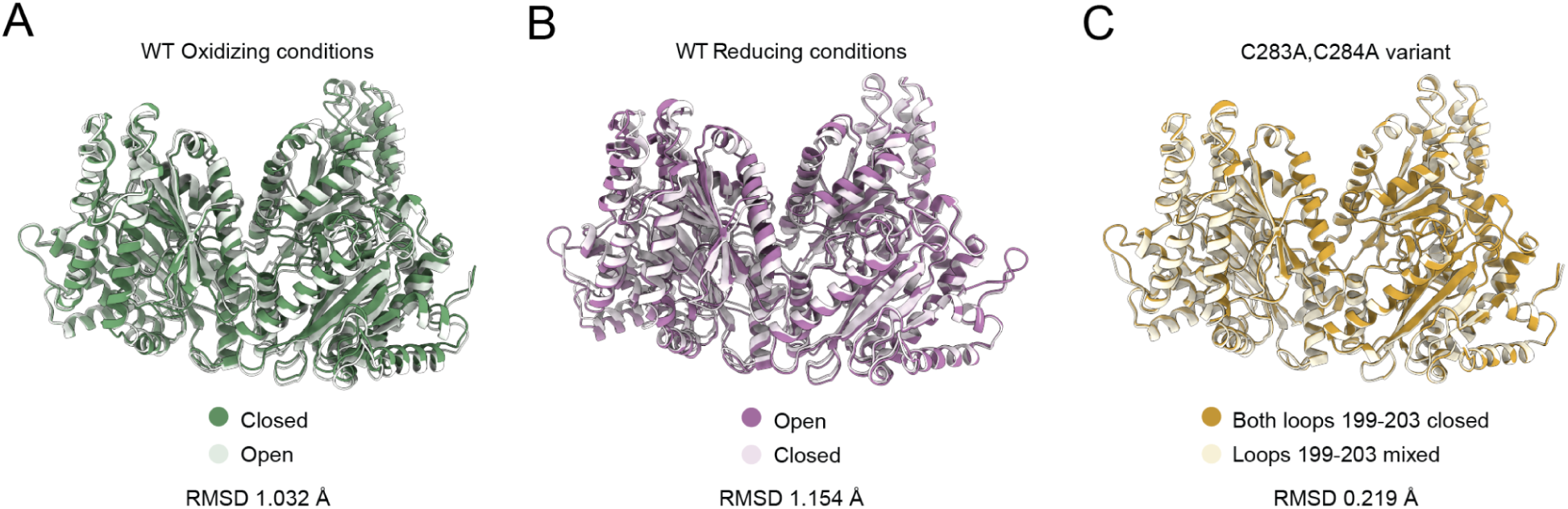
Superimpositions of the different dimer conformations found within each dataset. The major conformations found in each dataset are shown in darker shades, and the minor conformations in lighter shades. (A) Oxidized-closed vs oxidized-open. (B) Reduced-open vs reduced-closed. (C) C283A,C284A-open with loop 199-203 closed in both protomers (the one described in the results section) vs loop 199-203 in different states in the two protomers.

**SI Figure 9:**
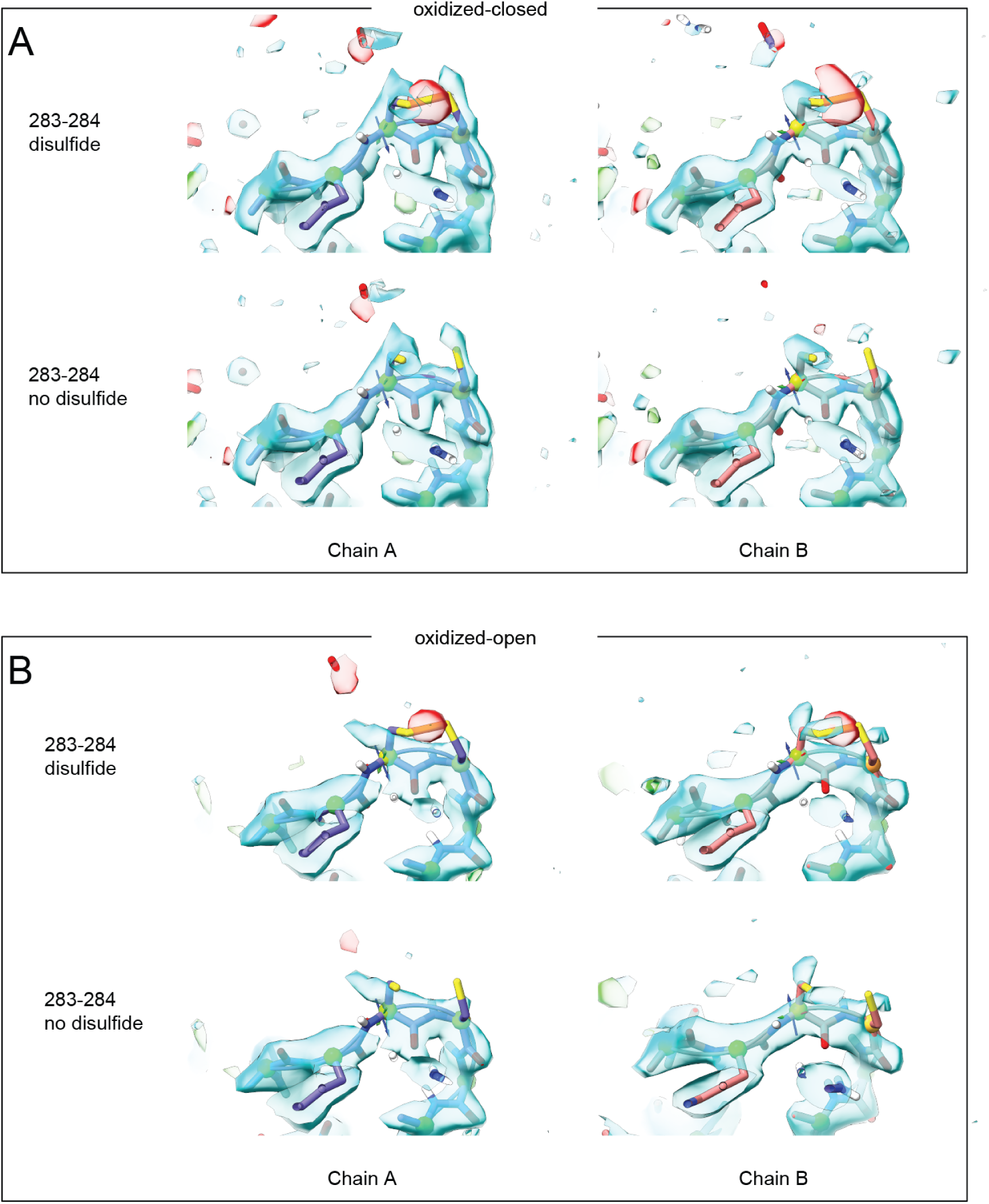
Modeling of the Cys283-Cys284 disulfide. Atomic models with or without a disulfide modeled between Cys283 and Cy284 were refined against the maps of the oxidized-closed (A) and oxidized-open (B) structures. The sharpened map (Fobs) from servalcat is shown as a translucent blue surface at a contour level of 2 sigmas. The difference map (Fobs-Fcalc) from servalcat is shown as translucent green/red surfaces at contour levels of +/- 5 sigmas for positive and negative difference peaks, respectively. Negative difference peaks (red) indicate that the data reject features of the model, while positive difference peaks (green) indicate a feature supported by the data but not present in the model.

**SI Figure 10:**
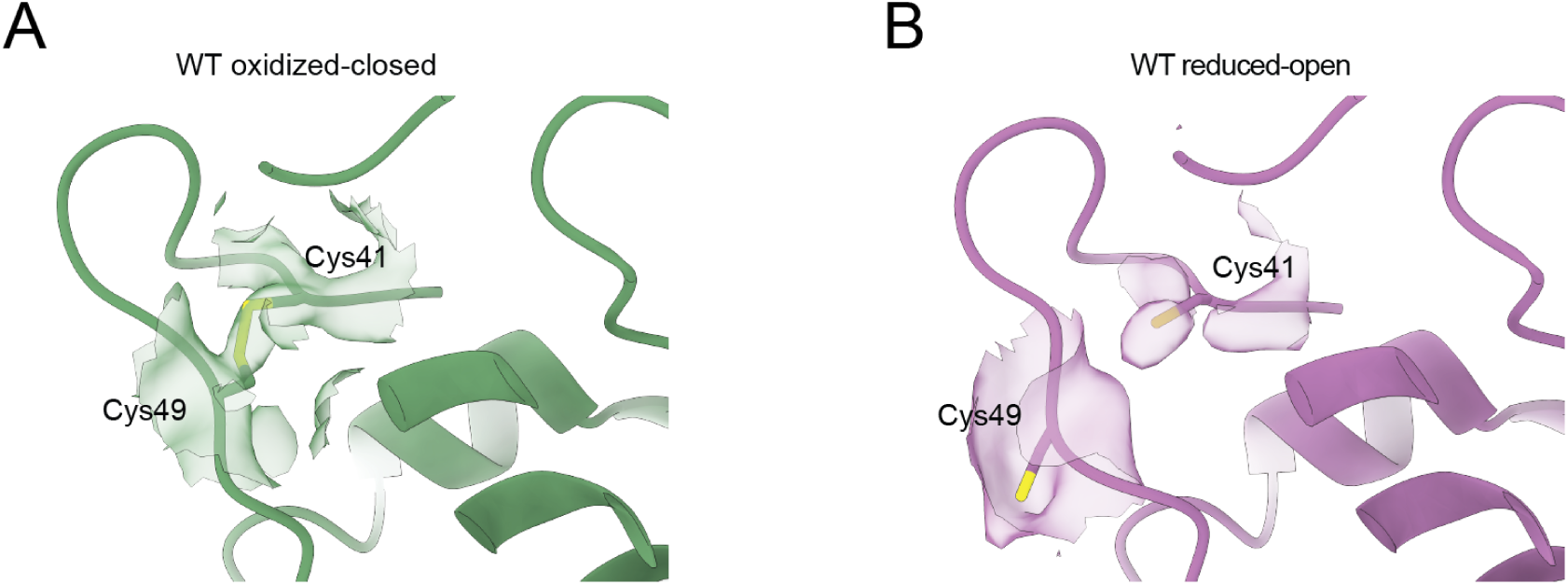
Comparison of the N-terminus between the oxidized and reduced structures. The non-regulatory Cys41-Cys49 disulfide reports on the redox state of the enzyme in the structures from different conditions. (A) In the oxidized-closed structure a disulfide is supported. (B) In the reduced-open structure no disulfide is observed and the cysteines are in fact not within bonding distance due to a different conformation of the flexible N-terminus. Unsharpened maps are shown as translucent surfaces around Cys41 and Cys49, at a contour level of 0.02.

### SI Movies

SI Movie 1: First component of variability of *Hn*CsoSCA under oxidizing conditions.

SI Movie 2: First component of variability of *Hn*CsoSCA under reducing conditions.

SI Movie 3: First component of variability of *Hn*CsoSCA C283A,C284A under oxidizing conditions.

---

**Table S1:** Cryo-EM data collection, refinement and model building statistics.

|  |  |  |  |  |  |  |  |  |  |
| --- | --- | --- | --- | --- | --- | --- | --- | --- | --- |
| Our internal ID | NV12 P410-J35 | NV12 P410-J56 | NV12 P410-J55 | NV21 P411-J26 | NV21 P411-J58 | NV21 P411-J48 | NV14 P445-J40 | NV14 P445-J61 | NV14 P445-J65 |
| Description | Ox, hexamer | Ox, dimer major | Ox, dimer minor | Red, hexamer | Red, dimer major | Red, dimer minor | Mut, hexamer | Mut, state 1 (both closed) | Mut, state 2 (closed, mixed) |
| Conditions | Oxidizing | Oxidizing | Oxidizing | Reducing | Reducing | Reducing | Oxidizing | Oxidizing | Oxidizing |
| Structure | oxidized-hexamer | oxidized-closed | oxidized-open | reduced-hexamer | reduced-open | reduced-closed | mutant-hexamer | mutant-open | mutant-open |
| EMDB entry | EMD-54972 | EMD-54973 | EMD-54974 | EMD-54975 | EMD-54976 | EMD-54977 | EMD-54978 | EMD-54979 | EMD-54980 |
| PDB entry | 9SKR | 9SKS | 9SKT | 9SKU | 9SKV | 9SKW | 9SKX | 9SKY | 9SKZ |
| Data collection and image processing |  |  |  |  |  |  |  |  |  |
| Microscope | Krios | Krios | Krios | Krios | Krios | Krios | Krios | Krios | Krios |
| Detector | K3 Bioquantum | K3 Bioquantum | K3 Bioquantum | K3 Bioquantum | K3 Bioquantum | K3 Bioquantum | K3 Bioquantum | K3 Bioquantum | K3 Bioquantum |
| Acceleration voltage (kV) | 300 | 300 | 300 | 300 | 300 | 300 | 300 | 300 | 300 |
| Nominal magnification | 130000 | 130000 | 130000 | 130000 | 130000 | 130000 | 130000 | 130000 | 130000 |
| Pixel size (Å/px) | 0.65 | 0.65 | 0.65 | 0.65 | 0.65 | 0.65 | 0.65 | 0.65 | 0.65 |
| Electron exposure (e−/Å2) | 64.76 | 64.76 | 64.76 | 64.76 | 64.76 | 64.76 | 68.90 | 68.90 | 68.90 |
| Nominal defocus range (µm) | -0.6 to -1.8 | -0.6 to -1.8 | -0.6 to -1.8 | -0.6 to -1.8 | -0.6 to -1.8 | -0.6 to -1.8 | -0.6 to -1.8 | -0.6 to -1.8 | -0.6 to -1.8 |
| Symmetry imposed | D3 | C1 | C1 | D3 | C1 | C1 | D3 | C1 | C1 |
| Initial particle images (no.) | 917605 | 1459839 | 1459839 | 818243 | 1635861 | 1635861 | 497507 | 1068384 | 1068384 |
| Final particle images (no.) | 486613 | 574591 | 277721 | 545287 | 543321 | 200733 | 356128 | 482720 | 525107 |
| Map resolution (Å) | 2.13 | 2.15 | 2.45 | 2.06 | 2.12 | 2.27 | 2.08 | 2.18 | 2.22 |
| FSC threshold | 0.143 | 0.143 | 0.143 | 0.143 | 0.143 | 0.143 | 0.143 | 0.143 | 0.143 |
| Map resolution range (min/median/max, Å) | 1.41 / 2.26 / 25.35 | 1.41 / 2.52 / 27.83 | 1.55 / 2.80 / 33.40 | 1.41 / 2.19 / 18.34 | 1.41 / 2.41 / 27.43 | 1.40 / 2.58 / 30.44 | 1.41 / 2.20 / 23.34 | 1.41 / 2.49 / 27.83 | 1.40 / 2.52 / 30.64 |
| Map sharpening B factor (Å2) | -73.7 | -58.4 | -71.7 | -72.6 | -59.1 | -58.5 | -72.7 | -63.4 | -65.7 |
| Atomic model building |  |  |  |  |  |  |  |  |  |
| Initial model used (PDB code) | AlphaFold2 prediction | AlphaFold2 prediction | AlphaFold2 prediction | AlphaFold2 prediction | AlphaFold2 prediction | AlphaFold2 prediction | AlphaFold2 prediction | AlphaFold2 prediction | AlphaFold2 prediction |
| Model-to-map resolution (masked, Å) | 2.23 | 2.21 | 2.55 | 2.20 | 2.23 | 2.44 | 2.14 | 2.25 | 2.31 |
| Model-to-map FSC threshold | 0.5 | 0.5 | 0.5 | 0.5 | 0.5 | 0.5 | 0.5 | 0.5 | 0.5 |
| Model composition |  |  |  |  |  |  |  |  |  |
| Non-hydrogen atoms | 7703 | 7764 | 7550 | 7740 | 7862 | 7627 | 7652 | 7652 | 7615 |
| Protein residues | 934 | 942 | 942 | 934 | 950 | 934 | 942 | 942 | 934 |
| Water molecules | 361 | 361 | 147 | 398 | 398 | 285 | 253 | 253 | 277 |
| Zn2+ ions | 2 | 2 | 2 | 2 | 2 | 2 | 2 | 2 | 2 |
| Model validation |  |  |  |  |  |  |  |  |  |
| Bond lengths RMSD (Å) | 0.012 | 0.013 | 0.012 | 0.011 | 0.012 | 0.012 | 0.012 | 0.012 | 0.013 |
| Bond angles RMSD (°) | 1.904 | 2.018 | 2.133 | 1.896 | 2.020 | 1.973 | 1.939 | 2.031 | 2.041 |
| MolProbity score | 1.81 | 1.37 | 2.47 | 2.05 | 1.98 | 2.08 | 1.81 | 1.47 | 1.88 |
| All-atom clashscore | 5.60 | 3.41 | 8.39 | 5.23 | 4.80 | 4.19 | 5.18 | 3.34 | 5.23 |
| Rotamer outliers (%) | 1.28 | 1.40 | 4.83 | 3.46 | 3.65 | 4.87 | 1.92 | 2.30 | 3.22 |
| Ramachandran favored (%) | 93.55 | 97.33 | 91.79 | 94.73 | 95.67 | 94.84 | 95.42 | 97.65 | 96.67 |
| Ramachandran allowed (%) | 6.02 | 2.56 | 7.25 | 4.95 | 3.59 | 4.41 | 4.58 | 2.35 | 3.01 |
| Ramachandran outliers (%) | 0.43 | 0.11 | 0.96 | 0.32 | 0.74 | 0.75 | 0.00 | 0.00 | 0.32 |
| Atomic displacement parameters (min/median/max, Å2) | 44.80 / 80.70 / 220.20 | 26.80 / 59.50 / 212.40 | 35.10 / 74.00 / 228.00 | 44.90 / 84.10 / 215.50 | 33.70 / 64.60 / 206.30 | 23.40 / 61.30 / 218.80 | 46.10 / 80.00 / 219.40 | 34.60 / 67.10 / 198.60 | 35.60 / 70.30 / 215.60 |
| Model to map fit |  |  |  |  |  |  |  |  |  |
| CC volume | 0.8906 | 0.9279 | 0.9066 | 0.8866 | 0.9177 | 0.9198 | 0.8988 | 0.9182 | 0.9131 |
| CC mask | 0.8933 | 0.9319 | 0.9109 | 0.8893 | 0.9212 | 0.9243 | 0.9011 | 0.9214 | 0.9158 |
| CC peaks | 0.4916 | 0.7505 | 0.7166 | 0.4791 | 0.7488 | 0.7469 | 0.4916 | 0.7295 | 0.7183 |
| Q-scores (min/median/max) | -0.791 / 0.870 / 0.963 | -0.872 / 0.887 / 0.963 | -0.685 / 0.834 / 0.938 | -0.770 / 0.870 / 0.966 | -0.758 / 0.879 / 0.954 | -0.759 / 0.858 / 0.958 | -0.882 / 0.882 / 0.970 | -0.782 / 0.876 / 0.957 | -0.785 / 0.869 / 0.950 |

